# Patient-derived triple negative breast cancer organoids provide robust model systems that recapitulate tumor intrinsic characteristics

**DOI:** 10.1101/2021.08.09.455691

**Authors:** Sonam Bhatia, Melissa Kramer, Suzanne Russo, Payal Naik, Gayatri Arun, Kyle Brophy, Peter Andrews, Cheng Fan, Charles M. Perou, Jonathan Preall, Taehoon Ha, Dennis Plenker, David A. Tuveson, Arvind Rishi, J Erby Wilkinson, W. Richard McCombie, Karen Kostroff, David L. Spector

## Abstract

Triple negative breast cancer (TNBC) is an aggressive form of breast cancer with poor patient outcomes, and an unmet clinical need for targeted therapies and better model systems. Here, we developed and comprehensively characterized a diverse biobank of normal and breast cancer patient-derived organoids (PDOs) with a focus on TNBCs. PDOs recapitulated patient tumor intrinsic properties and a subset of PDOs can be propagated for long-term culture (LT-TNBCs). Single cell profiling of PDOs identified cell types and gene candidates affiliated with different aspects of cancer progression. The LT-TNBC organoids exhibit signatures of aggressive MYC-driven basal-like breast cancers and are largely comprised of luminal progenitor (LP)-like cells. The TNBC LP-like cells are distinct from normal LPs and exhibit hyperactivation of NOTCH and MYC signaling. Overall, our study validates TNBC PDOs as robust models for understanding breast cancer biology and progression, paving the way for personalized medicine and tailored treatment options.

**Statement of Significance:** A comprehensive analysis of TNBC patient-derived organoids is presented by genomic, transcriptomic, and *in-vivo* analyses, providing insights into cellular heterogeneity and mechanisms of tumorigenesis at the single cell level.

## Introduction

Breast cancer is characterized into several histopathological subtypes based on the expression of various receptors: estrogen (ER), progesterone (PR) and human epidermal growth factor receptor 2 (HER2/ERBB2). Molecular subtyping revealed multiple subgroups based on gene expression patterns that have been refined overtime and constitute luminal A, luminal B, HER2-enriched, basal-like, and normal-like breast cancers (1–3). Luminal A (ER+/PR+/Her2-) is the most prevalent and constitutes ∼70% of all breast cancer cases, luminal B (ER+/PR+/HER2+) make up ∼12% of cases, HER2-amplified (ER-/PR-/HER2+) represents ∼5% of total cases and the basal or triple negative breast cancer (TNBC, ER-/PR-/HER2-) makes up ∼12% of total breast cancer cases (4). The majority of luminal breast cancers exhibit low proliferation rates while the TNBCs are highly aggressive and proliferative in nature. Since TNBCs lack ER/PR and HER2 receptors, patients are exempt from any targeted endocrine or HER2 therapies resulting in non-specific cytotoxic chemotherapy as the standard of care. The majority, but not all, TNBCs fall under the basal-like subgroup of breast cancer by gene expression based PAM50 profiling and a small subset belong to the highly invasive claudin-low subgroup (1,4–6). Further molecular characterization of TNBCs revealed heterogeneity within this subset and at least 6 molecular subtypes have been identified that can be used to design more targeted therapies (7, 8). TNBCs are typically presented as high-grade carcinomas (9) and show a significantly poor 5-year prognosis compared to other subtypes (10–12). Therefore, there is an unmet clinical need to better understand triple-negative disease progression and identify more precise druggable targets, and specific therapies for this cohort of breast cancer patients.

3-D organoid models have gained traction over recent years as the next generation of models for studying disease and development (reviewed in (13, 14)). While cell lines, spheroids and mouse model-derived organoids have been the primary *in vitro* model systems for studying cancer biology; patient-derived organoid (PDO) models have now been developed for a variety of cancers including those originating in the colon (15), pancreas (16), ovary (17, 18), prostate (19) and breast (20). Sachs et al. developed a methodology for growing human breast tumors *ex vivo* as organoid models that has now been expanded to the growth of organoids from normal mammary tissue as well (21, 22). While cancer cell lines have been a valuable resource for providing insights into cancer biology and drug development (23–25), PDO models are not only patient-specific but also provide a three-dimensional context which is closer to that of the actual tumor microenvironment. PDOs, therefore, represent unique model systems to study disease progression and for the identification and validation of better treatment options. However, we do not yet fully understand the extent to which these models recapitulate the cellular heterogeneity and complexities of triple negative disease.

Here, we developed a diverse breast cancer PDO biobank and performed comprehensive genomic, transcriptomic and cellular characterization of organoids with an emphasis on TNBCs. Using genomic assays we show that our organoid models recapitulate pathogenic single nucleotide variants (SNVs) and copy number alterations (CNAs) of breast cancers as portrayed in large scale breast cancer genomic datasets (10,26–28) and reveal lesser studied cancer driver genes. Transcriptomically, our biobank recapitulates the various subtypes and signatures of breast cancers, with a subset of organoids exhibiting signature profiles associated with poor patient outcomes. *In-vivo* transplants of these organoids highly recapitulated the patient-tumor morphology, providing strong evidence of retention of individual tumor intrinsic properties in long-term organoid cultures, even for models derived from rare BCs. In addition, we find that while normal PDOs retain the major cell types found within the mammary epithelium, the TNBC PDOs have lost this lineage specificity and are predominantly enriched for luminal progenitor (LP)-like cells. Single cell RNA-sequencing (scRNA-seq) of TNBC and normal PDOs identified differential signatures between the tumor and normal LP cells, providing insights into putative mechanisms of tumorigenesis. Lastly, we identified cells with various gene expression signatures in TNBC organoids that can be used to model and perturb various aspects of cancer biology, including tumorigenesis, hypoxia response, and EMT. Overall, our comprehensive characterization of TNBC organoids identified them as valid cancer models for studying cancer biology and for applications in precision medicine.

## Materials and Methods

### Patient Material

Tumor resections from breast cancer patients along with the distal and adjacent normal tissue were obtained from Northwell Health in accordance with Institutional Review Board protocol IRB-03-012 and IRB 20-0150. Specific information for all samples is available in Table S1. The collection of genomic and phenotypic data was consistent with 45 CFR Part 46 (Protection of Human Subjects) and the NIH Genomic Data Sharing (GDS) Policy. Informed consent ensured that the de-identified materials collected, the models created, and data generated from them can be shared without exceptions with researchers in the scientific community.

#### Patient-derived organoid culture

Patient-derived organoids were established and propagated using a previously published protocol (20). In summary, the tissues were manually cut into smaller pieces and treated with 2mg/ml collagenase IV in base media (ADF+++: Advanced DMEM-F12 (Invitrogen 12634-034) with 1x Glutamax (Invitrogen 12634-034), 10mM Hepes (Invitrogen 15630-056), 100U/ml Pen-Strep (Invitrogen 15140-122)) at 37°C for 45-90mins with gentle agitation to break the tissue into small clusters of cells. The suspension was intermittently resuspended by pipetting multiple times to ensure proper digestion of the tissue. The cell suspension was centrifuged at 300g for 5mins and the pellet was treated with red blood cell lysis buffer (Cat # 11814389001, Sigma) for 5mins at room temperature if it appeared bloody. The suspension was washed 2x with ADF+++ and plated in a matrigel (lot test for concentration of 8-10mg/ml, Cat # 356231, Corning) dome on pre-warmed tissue culture plates. The dome was incubated at 37°C for 15mins and supplemented with completed medium: 10% R-Spondin1 conditioned medium, 5nM Neuregulin 1 (Peprotech 100-03), 5ng/ml FGF7 (Peprotech 100-19), 20ng/ml FGF10 (Peprotech 100-26), 5ng/ml EGF (Peprotech AF-100-15), 100ng/ml Noggin (Peprotech 120-10C), 500nM A83-01 (Tocris 2939), 5uM Y-27632 (Abmole Y-27632), 1.2uM SB202190 (Sigma S7067), 1x B27 supplement (Gibco 17504-44), 1.25mM N-Acetylcysteine (Sigma A9165), 5mM Nicotinamide (Sigma N0636), 50ug/ml Primocin (Invitrogen ant-pm-1) in ADF+++. The organoids were passaged every 15-30 days using TrypLE^TM^ (Thermo Fischer 12605028) to break down the organoids into smaller clusters of cells and re-plating them in matrigel domes as described above. For tumor scrapings, the tumor surface was shaved on multiple sides to collect material which was subsequently manually broken down, treated with red blood cell lysis buffer and seeded in matrigel followed by regular PDO culture.

Organoid models labelled with the prefix HCM-CSHL were acquired as part of the Human Cancer Model Initiative (HCMI) https://ocg.cancer.gov/programs/HCMI and a subset of those models are or will be available for access from ATCC. The data for these models can be accessed here: dbGaP accession number phs001486. Organoid nomenclature: prefixes LNS, NH, DS, HCM-CSHL are de-identified patient IDs and are not distinct in any features other than protocols used for sample acquisition; prefix NM designates true normal samples collected from patients undergoing reductive mammoplasty (NM: <u>N</u>ormal <u>M</u>ammoplasty); suffixes: T=tumor, N=normal, ND=normal distal, NAdj=normal adjacent, and TSc= tumor scraping. Organoid freeze thaws are indicated in parenthesis: (passage frozen down)passage after thaw eg. NH85TSc (p4)p4.

#### Organoid DNA and RNA extraction

Organoid RNA was extracted using TRIzol® (Thermo Fischer 15596018) RNA extraction protocol. DNA was extracted by removing matrigel from organoids using ice cold PBS or TrypLE following by DNA extraction using Qiagen DNeasy Blood and Tissue kit (Qiagen 69504) with elution in nuclease free water (Thermo Fischer/Ambion 4387936).

#### Targeted gene panel sequencing

We performed capture based targeted gene panel sequencing (65) for a panel of potential cancer driver genes. Briefly, we used a panel of 143 cancer genes with a total of ∼4000 probes for capture. The captured DNA was paired-end sequenced with 150bp reads and a coverage of about 300-500x. Library preparation and sequencing of the targeted gene panel was performed by the CSHL Next Generation Sequencing Core Facility. We developed an analysis pipeline to prioritize identification of driver mutations. The sequencing reads are aligned to the hg19 reference genome using BWA (66), followed by conversion to BAM format and sorting with Samtools (67), removal of PCR duplicates with Picard (https://broadinstitute.github.io/picard/), and filtering with Bamtools (68) for mapping quality and proper read pairing. Coverage of the target regions is assessed for breadth and depth using Picard HSmetrics to ensure adequate coverage for confident variant detection. Variants are then called using VarScan2 (69) in somatic mode to stratify germline versus somatic variants, and are annotated with Annovar (70) to cover a broad range of variant assessment tools. We then select rare loss of function variants (nonsense, frameshift, splice site) with frequency less than 1% in the Gnomad, ExAC, EVS and 1000 Genomes databases. Missense and in-frame indel variants are selected if they are noted as pathogenic by ClinVar (71), or if they are both rare (<1% in all genome databases) and annotated as pathogenic by COSMIC (72), or if they are both rare and found to be present in the TCGA cohorts. Finally, missense variants are selected if they are annotated as potentially deleterious by the ensemble tools REVEL (73) and MCAP (74). Variants that were deleterious by REVEL and MCAP but did not have population level data were discarded from the final oncoplot. Oncoplots are generated from these candidate variants using Maftools (75).

#### RNA-seq

All RNA samples were quality controlled using a nanodrop followed by a bioanalyzer (RNA nano-kit Cat # 5067-1511) and only samples with RIN >7 were used for RNA sequencing. 750ng of RNA was used to prepared un-stranded RNA-seq libraries using Illumina TruSeq RNA Library prep kit v2 (RS-122-2001) and sequenced as 75bp paired-end reads.

#### RNA-seq analysis

The sequencing fastq files were quality checked using fastQC to make sure the reads were of consistent quality between different runs. The reads were aligned using STAR-aligner STAR-2.5.3a (76) using the following parameters:

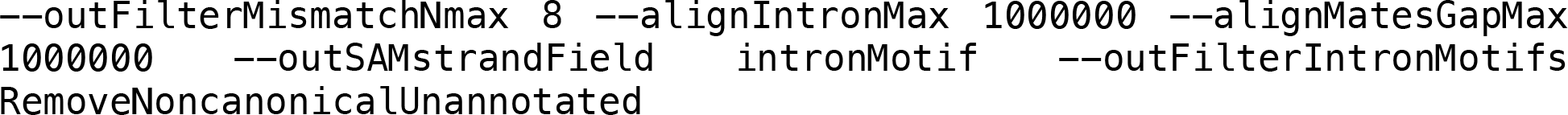

against the gencode v27 gtf reference file. Any PCR duplicates were marked in the aligned files using STAR with the following parameters:

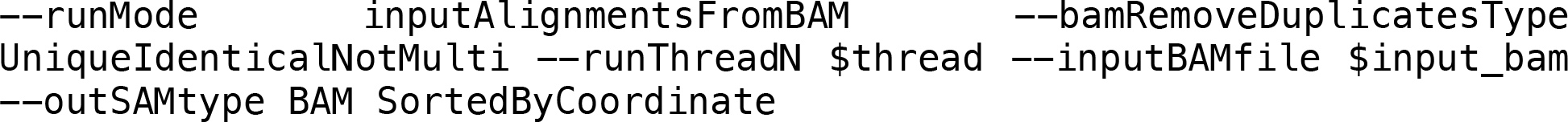

followed by removing duplicated reads using `smtools view -bF 0×400`. HTseq-count was used to count the reads per gene using the gencode v27 gtf file. The counts files were exported into R 4.1.0 and analyzed for differential expression using DeSeq2 1.32.0 (77). Concordance between technical replicates was ensured using PCA and sample distance matrix before summing them together for downstream analysis. Sample distance matrix was generated using euclidean distances between the samples and hierarchical clustering was performed using “ward.D2” linkage method followed by cutree with kmeans=6 with the R package “stats” (v4.1.0). For signature correlation, all the samples were used and an unsupervised clustering was performed for 838 previously curated gene expression signatures (37–39) and visualized using Java TreeView v1.2.0. For individual signature comparisons signature scores in experimental groups were compared using Kruskal-Wallis test followed by pairwise comparisons using Wilcoxon rank-sum test. Family-wise error rate was adjusted using Bonferroni-Holm method.

#### SMASH copy number

SMASH was performed as per the published protocol (32) starting with 750ng of genomic DNA. SMASH was performed in batches of 10 samples and sequenced on a MiSeq PE150bp run. The SMASH analysis is based on identification of Maximal Unique Matches (MUMs) to the human genome in all read pairs (32). These MUMs were filtered to remove matches <20 bp, matches with <4 bp of excess unique sequence, and matches on read 2 that are within 1000 bases of the matches from read 1. Raw copy number profiles are then generated from the remaining 3–4 matches per read pair which are then added to empirically sized bins spanning the genome. Regions with identical copy are expected to yield similar bin counts using these empirical bins. We next perform GC correction by normalizing counts based on LOWESS smoothing of count vs. GC content data in each bin. Final copy number profiles are normalized so that the autosome has an average copy number of 2. Plots were generated with G-Graph MUMdex software (https://mumdex.com/) and IGV browser v2.9.2 (78).

#### Organoid Histology

Organoid domes in complete medium were scraped from the tissue culture plate and collected in falcon tube precoated with BSA. The organoids were collected and washed 1x with PBS by spinning at 300g for 5mins. Organoid harvesting solution (Cat # 3700-100-01, Trevigen) was added to the organoids (3x the volume of matrigel) and incubated at 4°C on ice for 30minutes to ensure that matrigel was removed and the organoids were concentrated at the bottom. The organoids were washed 1x with ample PBS and fixed with fresh 4% PFA at room temperature for 10minutes. 1:1 (v/v) BSA was added to the tube and spun at 300g for 5mins to remove the PFA. The organoids were washed 2X with ample amounts of PBS and embedded in 2% agarose in dH2O). The agarose organoid molds were then paraffin embedded and cut into 5um sections.

#### Organoid Hematoxylin & Eosin (H&E) staining and Immunohistochemistry (IHC)

H&E and IHC staining were performed at the CSHL Histology Core Facility. PFA fixed organoids in agarose were processed in Thermo Excelsior ES processor and embedded with Thermo HistoStar embedding system following manufacturer’s protocols. Paraffin blocks were cut into 5um sections and mounted onto positively charged slides (VWR superfrost plus micro slide).

For H&E staining, slides were stained in a Leica Multistainer (ST5020). Briefly, slides were deparaffinized and rehydrated and then stained in hematoxylin (Hematoxylin 560 MX, Leica) for 1 min, followed by destaining in Define MX-aq (Leica) for 30 sec, bluing in Blue Buffer 8 (Leica) for 1min and subsequently stained in eosin (EOSIN 515 LT, Leica) for 30sec. After dehydration, coverslips were placed onto glass slides using a robotic coversliper (Leica CV5030).

IHC slides were stained in Discovery Ultra automatic IHC stainer (Roche) following standard protocols. Briefly, slides were subjected to antigen retrieval (Benchmark Ultra CC1, Roche) at 96°C for 1hr; primary antibodies were incubated at 37°C for 1hr and Discovery multimer detection system (Discovery OmniMap HRP, Discovery DAB and Purple, Roche) was used to detect and amplify immuno-signals. Antibodies used: Ki67 (Spring Bioscience, #M3062, 1:500).

#### Organoid formation assay

Organoids were processed using TrypLE and 1500 single cells per well of a 96 well plate were seeded in 10% matrigel + complete growth medium. Cell viability was assessed using Cell Titre Glo 3D luminescence assay (Promega G9683 CellTiter-Glo 3D Cell Viability Assay). Baseline cells were measured using Cell Titre Glo 3D assay at 24 hrs (d1) post seeding and growth was measured at 6 days after seeding (d6). Each organoid line was evaluated for multiple passages n=2 or n=3 per PDO.

For MYCi and DAPT experiments: Organoids were processed to single cells using TrypLE and seeded in 50ul matrigel domes as 10,000 cells per well of a 24 well-plate. Complete medium or medium supplemented with DMSO, DAPT (Selleckchem S2215 DAPT) and MYC-inhibitor (Selleckchem S8906 MYCi975) were added to the respective wells. For normal PDOs, organoids were dissociated with trypLE and live cells were sorted using fluorescence activated cell sorting (FACS) based on staining of CD46f, EPCAM and 7AAD. Organoids were allowed to form for 12 days, images were acquired using microscope and organoids were manually counted for each condition. Statistical analysis was performed using one-way ANOVA test followed by pairwise comparisons using two-sample t-test. Family-wise error rate was adjusted using Bonferroni-Hold method.

#### Organoid proliferation index analysis

Paraffin embedded organoid sections were IHC stained for Ki67. Slides were scanned and viewed using Aperio ImageScope 12.3.3. Images were analyzed in FIJI (79). Briefly, the images were deconvolved using Colour Deconvolution for hematoxylin and DAB, converted to 8-bit binary images and analyzed using the BioVoxxel Toolbox plugin (https://www.biovoxxel.de/#/) to evalute %Ki67 positive cells per organoid. Multiple passages for each organoid line were evaluated, n=2 or n=3 per PDO.

#### Drug dose response assays

Organoids were digested into single cells and seeded as 1500 cells/well of a 384 wp (USA-Scientific Cat # 5678-1976) as suspension cultures in complete medium with 10% matrigel using a liquid handler. Organoids were incubated at 37C for 24hrs and drugs were added using a drug dispenser Beckman Echo 650. The organoids were incubated for 5 days at 37C and assayed using the CellTiter-Glo® 3D Cell Viability Assay (Promega Cat # G9681) and read for luminescence using the EnVision 2105 plate reader. The data was analyzed using GraphPad Prism v9.0.0. The screens were performed in 3 technical replicates with 2-4 experimental replicates per organoid line. The raw reads were first normalized against untreated controls and a non-linear model was fit for the mean, after removing any detected outliers, using the log(inhibitor) vs normalized response with a variable slope.

#### Animals

Six-week-old female NOD scid mice (NOD.Cg-*Prkdc*^scid^/J) were obtained from the Jackson laboratory (JAX stock #001303) and acclimated at the Cold Spring Harbor Laboratory Animal Shared Resource for a minimum of 1week. All animal experiments were performed in accordance with the Institutional Animal Care and Use Committee.

#### *In-vivo* transplant experiments

TNBC organoids were harvested using the organoid harvesting solution (Trevigen Cat# 3700-100-01) and manually counted. Organoids were resuspended in a 1:1 mixture of PBS:matrigel and 50K organoids were injected into the bilateral mammary fat pads by the fourth nipple of female NOD scid mice (NOD.Cg-*Prkdc*^scid^/J). Mice were anesthetized with 1.5-2% isoflurane and weighed before the injections. The animals were regularly monitored for their weight, tumor size and any other discrepancies. Mice were sacrificed when the tumors reached any of the following end-points: 2cm tumors, ulceration, visible necrosis, blistering of tumors or deteriorating health of the mice. At end-point, dissections were performed and the tumors along with lungs, liver, lymph nodes and the femur were fixed in 4% PFA overnight at 4°C. If the tumors were not observed the mammary fat pads were collected instead. The transplant experiments were done with 2 independent passages of PDOs, with 4-6 injections per PDO per passage.

Fixed tumors and tissues were processed for histology as above. Metastases and micro-metastases were assessed using IHC with a human mitochondria antibody (Millipore MAB1273 Anti-Mitochondria clone 113-1).

#### Flow Cytometry

Organoids were scraped in the culture medium and washed 1X with PBS. TrypLE^TM^ was used to fully digest the organoids into single cells. The cells were counted, diluted to 200,000 cells/100ul and stained in 100ul of ADF+++ using anti-Epcam (1:50), anti-CD49f (1:50), 7AAD (1:50). The following antibodies were used: PE Mouse IgG2a, κ Isotype Control (BD 555574), APC Mouse IgG2b κ Isotype Control RUO (BD 555745), Alexa Fluor® 647 anti-human CD326 (EpCAM) Antibody (Biolegend 324212), PE anti-human/mouse CD49f Antibody (Biolegend 313612). The cells were read using a BD Fortessa and analyzed using the FACS DIVA and FlowJo v10 software.

#### Single cell RNA-seq

Organoids were digested into single cells using TrypLE^TM^, resuspended in 0.04% BSA in PBS as 1 million cells/ml. ∼12,000 cells were loaded into one well of a 10x Chromium microfluidics chip. Single cell barcoding and libraries were prepared using the 10x Chromium v3 chemistry (Cat # V3 reagents 1000075, V3 chips -1000153 or NextGEM reagents-1000121, chips 1000120). Libraries were quality checked using a Bioanalyzer HS kit for cDNA yield and final library size and qubit to quantify.

Single cell analysis was performed in three different batches (Table S7). Batch 3 was a multiplexed pool of 4 samples, which were demultipexed using a custom genotype-aware pipeline. At the time of 10X Genomics library preparation, ∼20,000 cells from each of the four organoids were set aside to prepare low-input bulk RNA-seq libraries tagged with unique i7 barcodes. These bulk libraries share the same adapter structure as 10X Genomics libraries, and were spiked into the Illumina NextSeq500 flow cell at a 5% molar ratio to obtain ∼5M reads per organoid. These barcoded bulk libraries were then used to create reference VCF files using cellSNP v0.3.2 by searching a list of 7.4M common human SNPs from the 1000 Genomes Project (http://ufpr.dl.sourceforge.net/project/cellsnp/SNPlist/genome1K.phase3.SNP_AF5e2.chr1toX.hg38.vcf.gz). Genotype profiles were filtered to include only positions with < 10% minor allele frequency and >20 UMI counts. In parallel, per-cell VCF files were generated from the multiplexed single cell library using the cellSNP with the same parameters. Cells from the single-cell pool were assigned to their respective donors using Vireo v0.4.2 (80). *Genotyping low-input bulk RNAseq library prep:*

RT Primer Design:

CTACACGACGCTCTTCCGATCTSSSSSSSSNNNNNNNNNNVVVVVTTTTTTTTTTTTTTTTTTTTTTTTTTTTTTVN

where:

SSSSSSSS = 8bp sample barcode, with following multi-plexing key: GACAGTGC=HCM-CSHL-0366-C50

GAGTTAGC=NH85TSc

GATGAATC=NH95T GCCAAGAC=NH93T

NNNNNNNNNNVVVVV = 15bp UMI with 5 non-T residues at 3’ end Template Switch Oligo: AAGCAGTGGTATCAACGCAGAGTGAATrGrGrG cDNA_amplification_Forward:

AATGATACGGCGACCACCGAGATCTACACTCTTTCCCTACACGACGCTCTTCCG

cDNA_amplification_Reverse: AAGCAGTGGTATCAACGCAGAGT

RT was performed using the SuperScript IV (Life Technologies #18091050) according to the manufacturer’s instructions except for the addition of 1uM Template Switch Oligo during first strand synthesis. All custom oligos were synthesized by IDT. After cDNA amplification, molar concentration was estimated using the Agilent Bioanalyzer 2100, libraries were pooled at an equimolar ratio, and prepared for Illumina sequencing using the Nextera XT DNA Library Prep Kit (Illumina) according to the manufacturer’s instructions. Final fragmented libraries were again checked and quantified by Bioanalyzer prior to mixing at a 1:20 molar ratio with 10X Genomics libraries for sequencing. *Sequencing and Mapping:* The libraries were sequenced on Illumina NextSeq 500 High Output 75 cycle kits using the read format: 8bp (i7 index) x 28bp (Read 1) x 56bp (Read2). 10X Genomics libraries were mapped using Cell Ranger version 4.0.0 (10X Genomics) with default settings and a custom genome reference based on the comprehensive gene annotation set from Gencode Release 32 (GRCh38.p13) (http://ftp.ebi.ac.uk/pub/databases/gencode/Gencode_human/release_32/gencode.v32.annotation.gtf.gz). For the multiplexed pool, sample identities for each cell were assigned using Vireo as described above, such that each sample could be subset from the pooled matrix during analysis.

Filtering, feature selection, clustering, and other secondary analyses were carried out in R using Seurat v4.0.3 (https://satijalab.org/seurat/) (47, 81). Gene set enrichment analyses were performed using GSEA v4.1.0 (82). Following gene sets were used for the various signature scores: adult human breast epithelium markers from (49), Mammary epithelial lineage scores from (40), NOTCH signaling: REACTOME_SIGNALING_BY_NOTCH, BMP2 targets: LEE_BMP2_TARGETS_UP, MYC signature from (83), Hypoxia signature: HALLMARK_HYPOXIA, Basal mammary stem cell signature from (40).

#### Quantification and statistical analysis

Statistical analyses were performed using R (version 4.1.0) on RStudio and GraphPad Prism software (v9.1.2, GraphPad Software, San Diego, California USA, www.graphpad.com). Specific tests are indicated in the figure legends along with the statistical significance.

#### Materials, data and code availability

Organoid lines generated under the HCMI project (starting with HCM) will be available for purchase from the American Type Culture Collection (ATCC). All raw data will be available to download from dbGaP (phs002722.v1.p1 and phs001486). Copy number segment file is available in Table S2, transcriptome signature scores are in Table S4. Metadata for the different PDOs is available in Table S1. All original code will be deposited at https://github.com/bhatia-sonam/manuscript-v1

## Results

### Establishment and somatic variant profiling of a diverse patient-derived breast cancer organoid biobank

Breast cancer tissues along with paired normal breast tissues were acquired from female patients and developed into 3-D organoids. In addition to breast cancer tissues, we also collected normal reductive mammoplasty samples from 10 cancer-free patients to generate normal PDOs for downstream analyses. The majority of tumor samples were from patients with invasive ductal carcinomas (56/87, Fig. 1A), 11/87 were from invasive lobular carcinomas, 4/87 were from metastatic lymph nodes and a small percentage from other categories (Fig 1A, Table S1). The majority of the tumor samples (43/87) were luminal BC as defined by immunohistochemistry (ER/PR+/HER2-), 37/87 were from TNBCs (ER-PR-HER2-), 2/87 were from ER/PR+/HER2+ BCs, 2/87 were from ER-/PR-/HER2+ subtype, and 3/87 samples belonged to post-treatment residual tissue with no visible carcinoma (NA) (Fig 1B). Samples were collected from patients of various age groups (Fig 1C) and diverse ethnic and racial backgrounds (Fig 1D). An increased proportion of TNBC tumor samples was observed from black patients with African or west-Indian heritage (Fig 1D, Table S1) which recapitulates the higher incidence of TNBCs in the African-American community (29, 30). Using a DNA-seq panel of 143 cancer driver genes, we identified pathogenic single nucleotide variants (SNVs) in 49 tumor organoid lines (Fig 1E); *PIK3CA* was mutated in 33% of the tumor organoid lines, the majority of which were from patients with luminal breast cancer. 24% of the organoid lines (45% of TNBC PDOs 10/22) had a pathogenic *TP53* mutation and primarily represented the high grade TNBC cohort (Fig 1E). Two of the organoid lines are derived from a patient with rare breast cancer showing adenoid cystic carcinoma (AdCC) like morphology, NH87T (primary tumor) and HCM-CSHL-0655-C50 (lymph node metastasis from the same patient) (Table S1). The AdCC organoids show a non-traditional TNBC mutation profile with pathogenic mutations in *APC*, *KDM6A* and *NOTCH1* (Fig 1E), which were previously observed in AdCCs of the breast (31). Mutations in *KMT2C* and *GATA3* were also observed in a variety of luminal breast cancer organoids, *CDH1* mutations were present in 10% of PDOs all of which belonged to invasive lobular carcinomas, and *ARID1B* mutations were found in a few TNBC-derived organoids (Fig 1E). Overall, the mutation profiles of patient-derived BC organoids are in concordance with the mutational landscape of BCs (10, 26). A relatively large subset of tissues (24/87) that resulted in cultured organoids did not have pathogenic SNVs (Fig 1F, Table S1), while 10/87 samples dropped off in culture at early passages (Fig 1F). The PDOs with pathogenic SNVs showed a range of growth properties *in vitro* with subsets exhibiting long-term continued expansion while others seem to have more limited cultures either in terms of maximum passages reached or limited expansion abilities (Table S1).

**Figure 1.**
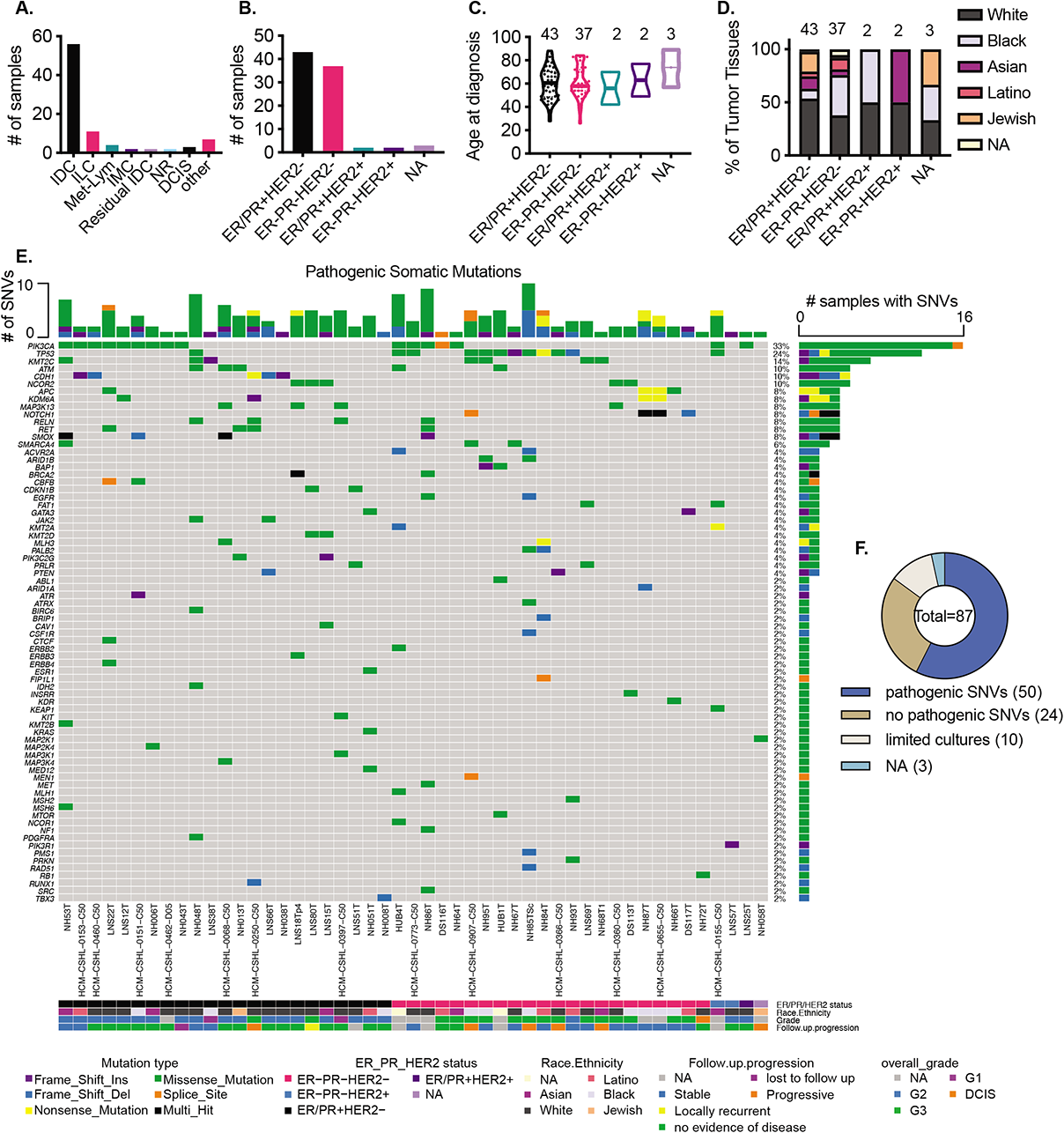
Establishment and somatic variant profiling of the breast cancer organoid biobank. **A.)** Summary of cancer type of the various tumor tissues that were used to generate organoids. IDC: Invasive ductal carcinoma, ILC: Invasive lobular carcinoma, Met-Lym: lymph node metastasis, IMC: Invasive mucinous carcinoma, NR: no residual tumor seen, DCIS: Ductal carcinoma in-situ, other: see Table S1 **B.)** Histopathological subtypes of the tumor tissues, ER/PR: Estrogen receptor (ER) and/or Progesterone receptor (PR), NA: not assessed **C.)** Age at diagnosis of the various subgroups of patient tumor tissues **D.)** Subtype specific, self-identified racial and ethnic breakdown of the patients represented in this biobank **E.)** Pathogenic single nucleotide variants (SNVs) identified in putative cancer driver genes in patient-derived organoids. **F.)** Proportion of organoids with pathogenic SNVs identified. Pathogenic SNVs: SNVs called pathogenic by ClinVar, COSMIC or REVEL and MCAP scores from targeted gene-panel sequence (49 samples) or whole exome sequencing (1 sample) (see Table S1), limited cultures: cultures where organoids were established at early passages (p0-p1) but could not be propagated, NA: not assessed

A comparison of the SNV profile between several pairs of patient tumor and PDOs showed high concordance of the pathogenic SNVs between the tumor and tumor derived organoids (Fig S1A), except in some cases where the SNVs are present at a much lower variant allele frequency (VAF) in the tumor, for instance, NH84TT-P0 showed <25% TP53 VAF compared to 100% VAF in NH84TT-p10 PDO (Fig S1A) suggesting normal contamination in the tumor tissues and a successful outgrowth of cancer cells in organoid cultures. Longitudinal analyses for some patient-derived organoids showed the stability of primary driver mutations overtime in culture (Fig S1B). To determine whether the amount of tumor material was a limiting step for generation of organoids, we collected tumor scrapings (see methods) from a subset of tumor samples. A comparison of tumor tissue (labeled TT) and the scrapings (labeled Sc) from three different patients (Fig S1C) showed high concordance and indicates that, if necessary, small amounts of tumor material can be used to generate organoids. We also examined the pathogenic SNVs in TNBC tissue samples (p0) that did not result in a successful generation of organoids (establishment or long-term cultures) and 6/10 had TP53 mutations and some samples also had BRCA1/2 mutations (Fig S1D), the latter were largely absent in successful organoid cultures (Fig 1E).

### Breast cancer organoids are enriched for copy number alterations

Having identified pathogenic driver mutations for the tumor-derived organoids, we performed copy number analysis using SMASH (short multiply aggregated sequence homologies) (32) for various PDOs. TP53 mutated TNBC PDOs make up the majority of the ER-subset and are highly genomically altered compared to the luminal ER+ organoids (Fig 2A, B, Table S2). Some organoids that were deemed to be derived from tumor based on pathogenic SNVs (eg. LNS12T, LNS18T, NH06T etc.) showed no prominent CNAs. Some TNBC PDOs that showed trace SNVs such as NH58T, NH72T and NH66T also showed minimal CNAs (Fig 1E and 2A). In keeping with large scale genomic datasets (10, 26), a consistent gain of chromosome 1q (chr1q) and loss of chr16q was observed in the luminal organoids where CNAs were detected (Fig 2A). The TP53 mutated TNBC organoids were highly genomically aberrant where multiple lines showed a gain of chromosomes (chrs) 1q, 8q, 19q and chrs 7, 20 and 21; loss of chrs 3p, 4, 5, 17p and Xp was also observed in multiple organoid lines (Fig 2A). These TNBC PDOs also showed copy number loss of chr4q and chr5q (Fig 2A) that have been previously reported to be over-represented in basal-like breast cancers (33). Of note, the AdCC-like organoids, NH87T (primary tumor) and HCM-CSHL-0655-C50 (lymph node metastasis from the same patient), were considerably less genomically aberrant as compared to the other TNBC organoids and showed focal amplification of chr8q and deletion of chr6p (Fig 2A, C-D). Interestingly, the chrX deletion was only observed in the primary tumor (NH87T) sample (Fig 2A, C) and a focal deletion was observed in chr2 of the lymph met HCM-CSHL-0655-C50 (Fig 2C). The AdCC organoid CNA profiles are consistent with the low CNA profiles that are typically observed in AdCC-like breast cancers (31).

**Figure 2.**
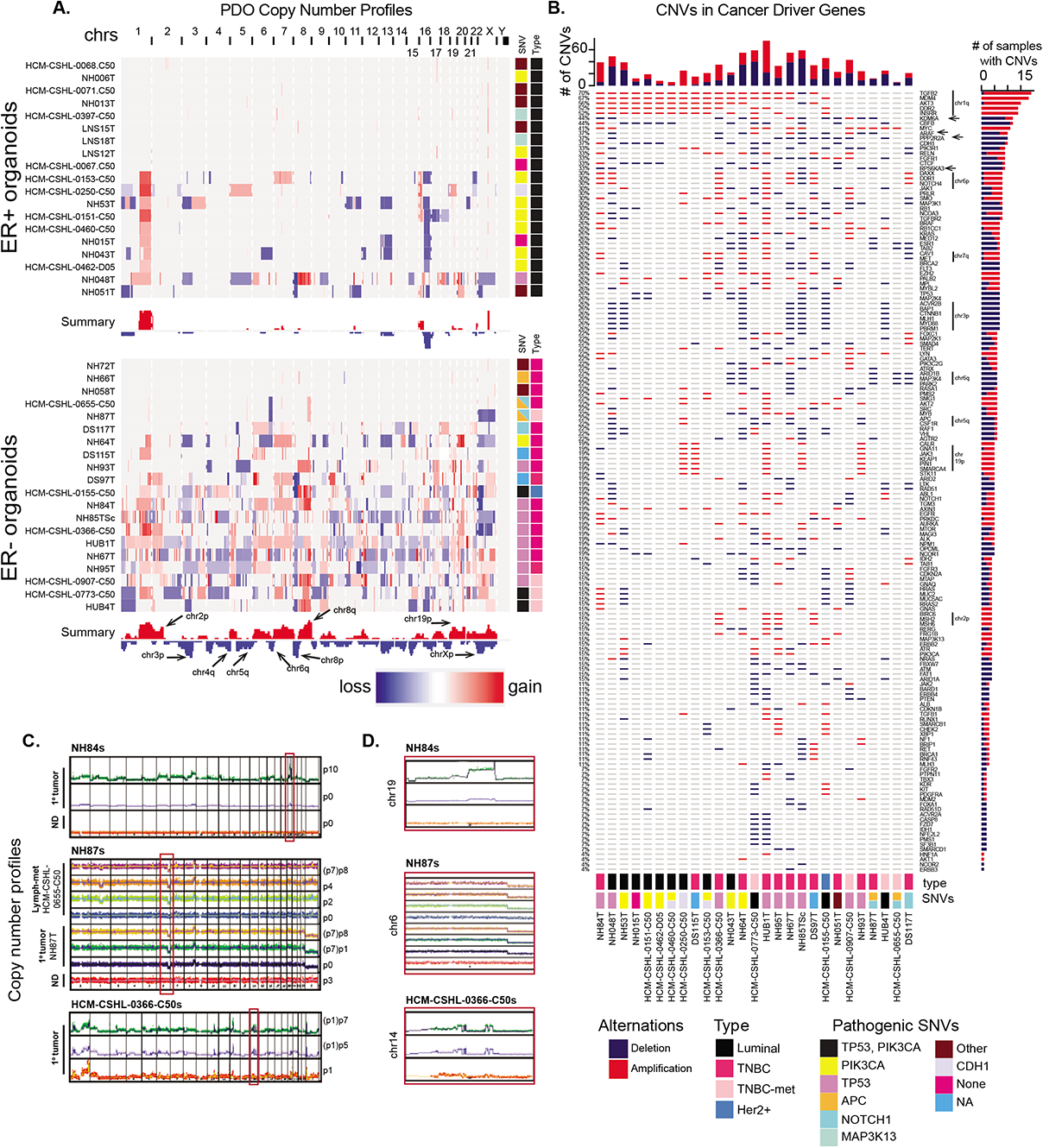
Copy number alterations (CNAs) enriched in the organoid models. **A.)** Copy number profiles, from IGV, of the various ER+ and ER-tumor derived organoids, along with the summary of overall copy number alterations across all samples. Side panel shows the pathogenic SNVs identified in that sample from Fig 1E. **B.)** Copy number amplifications or deletions identified in putative cancer driver genes (from Fig 1E). **C.)** Copy number across different passages of three different sets of patient-derived organoids. **D.)** Magnified view of the chromosome regions in the red boxes in C.

For a subset of these samples, we compared the copy number profiles of the primary tumor with the paired tumor and normal organoids and observed more pronounced CNAs in the organoids compared to the tumors suggesting that the organoid culture enriches for tumor cells (Fig 2C-D). As observed in the SNV data (Fig S1) the NH84T tumor (p0 with <25% VAF of pathogenic TP53) was likely very heterogenous and contained a significant population of normal cells, however, the tumor cells successfully outgrew the normal and resulted in a highly pure tumor organoid culture over multiple passages (Fig 2C-D). For HCM-CSHL-0366-C50 and NH87T the early passage p1 or tumor tissue (p0) respectively, had relatively less normal contamination and maintained their copy number profiles over time in culture (Fig 2C-D). While copy number profiles were enriched for in successful tumor PDOs, for some cultures we observed loss of copy number alterations in the organoids over-time, suggesting a normal outgrowth (Fig S1E).

Since tumor organoids are more enriched for tumor cells compared to the p0 tumor tissue, we profiled the putative cancer driver gene panel in individual organoid samples. TNBC PDOs typically had a higher copy number alteration frequency of these cancer driver genes (top bar-graph in Fig 2B). *TGFB2*, *MDM3*, *AKT3*, *DDR2* and *INSRR* showed the highest frequency of alteration (Fig 2B) and are all genes present on chr1q that is amplified in both luminal and TNBC PDOs (Fig 2A-B). Interestingly, we also found a higher frequency of deletion of *KDM6A*, *ARAF* and *RPS6KA3* tumor suppressors, located on chrXp, specifically in the TNBC-derived organoids (arrows in Fig 2B). Loss of *PPP2R2A* (chr8p), which was previously reported to be a tumor suppressor in breast cancer (10), was also identified in 38% of the samples, the majority of which are of TNBC subtype (Fig 2B). Furthermore, there is an over-representation of loss of chr3p in the TNBC samples, that results in the deletion of potential tumor suppressors: *ACVR2B*, *BAP1*, *CTNNB1*, *MLH1*, *MYF88* and *PBRM1* (Fig 2B). Copy number loss of chr6q is also common in TNBC organoids and results in loss of *ARID1B*, *MAP3K4*, and *PARK2*, a master regulator of G1/S cyclins (34). *ESR1*, the gene encoding estrogen receptor (ER), present on chr6q is also frequently lost in these TNBC organoids (Fig 2A-B). We also identified gains of *CALR*, *JAK3*, *KEAP1*, *PIN1* and *SMARCA4* that are associated with the amplification of chr19p and amplification of mismatch repair (MMR) genes *MSH2* and *MSH6* located on chr2p (Fig 2A-B) in subsets of TNBC PDOs. While MMR genes are commonly mutated in various cancers, their overexpression was recently associated with aggressive prostate cancers (35) and was shown to promote genomic instability in yeast (36) but their role is yet to be determined in TNBC disease progression.

Taken together, the SNV and copy number profiling of PDOs shows robust retention of genomic features of various types of breast cancers, including luminal, TNBC, and rare AdCC-like carcinomas. While there might be small alterations overtime in PDO cultures, the key pathogenic mutations and overall copy number profiles are conserved throughout organoid culture (Fig S1, 2C-D) supporting their utilization as valid cancer models. Additionally, we find overrepresentation of some lesser studied copy number variants in our data such as loss of tumor suppressors *RPS6KA3*, *PPP2R2A* and *PARK2* and copy number gains of MMR genes *MSH2* and *MSH6* that might have important unexplored consequences in breast cancer progression.

### A subset of TNBC organoids recapitulate signatures of aggressive basal-like breast cancers

Next, we performed RNA-seq on various tumor and normal-derived organoids to profile their transcriptomes (Table S3). Hierarchical clustering of samples by euclidean distance divided them into six groups (Fig S2A). We used supervised clustering with 838 previously curated gene expression signatures (37–39) (Table S4) and found similar patterns of clustering among the various organoid lines (Fig 3A). Group 4 is largely comprised of the true normal organoids (labeled with prefix NM in Fig S2A) derived from breast tissues obtained from individuals undergoing reductive mammoplasty (Normal in Fig 3A-B, S2A, C), paired normal PDOs derived from normal tissue adjacent or distal to the tumor and very few tumor-derived organoids; this cluster showed an enrichment of normal mammary stem cell (MaSC) signatures and low proliferation related expression profiles (Fig 3A-B). Group 1 is comprised of a mixture of either paired normal organoids or tumor organoids that did not have a strong driver mutation (eg. *PIK3CA* or *TP53*) (Fig 3A, S2A, 1E). This group had a signature profile that was distinct from the true normal group (Group 4) and is most similar to luminal cell signatures (Fig 3A). The samples in this group also have a higher proliferation signature compared to the true normal Group 4 (Fig 3A). Group 5 mostly contained PDOs derived from Luminal BC and showed luminal like gene-expression signatures (Fig 3A-B). Signature profiling also uncovered immune and stromal signatures that highly corresponded to samples in Group 3 (Fig 3A, S2C). Interestingly, two of the samples in this group NH63T and NH54T exhibited limited propagation in culture due to stromal outgrowth leading us to hypothesize that these samples had some fibroblast-like cells. We also found a strong interferon (IFN) signature in this group along with some additional PDOs from Group 6 (Fig 3A, S2C).

**Figure 3.**
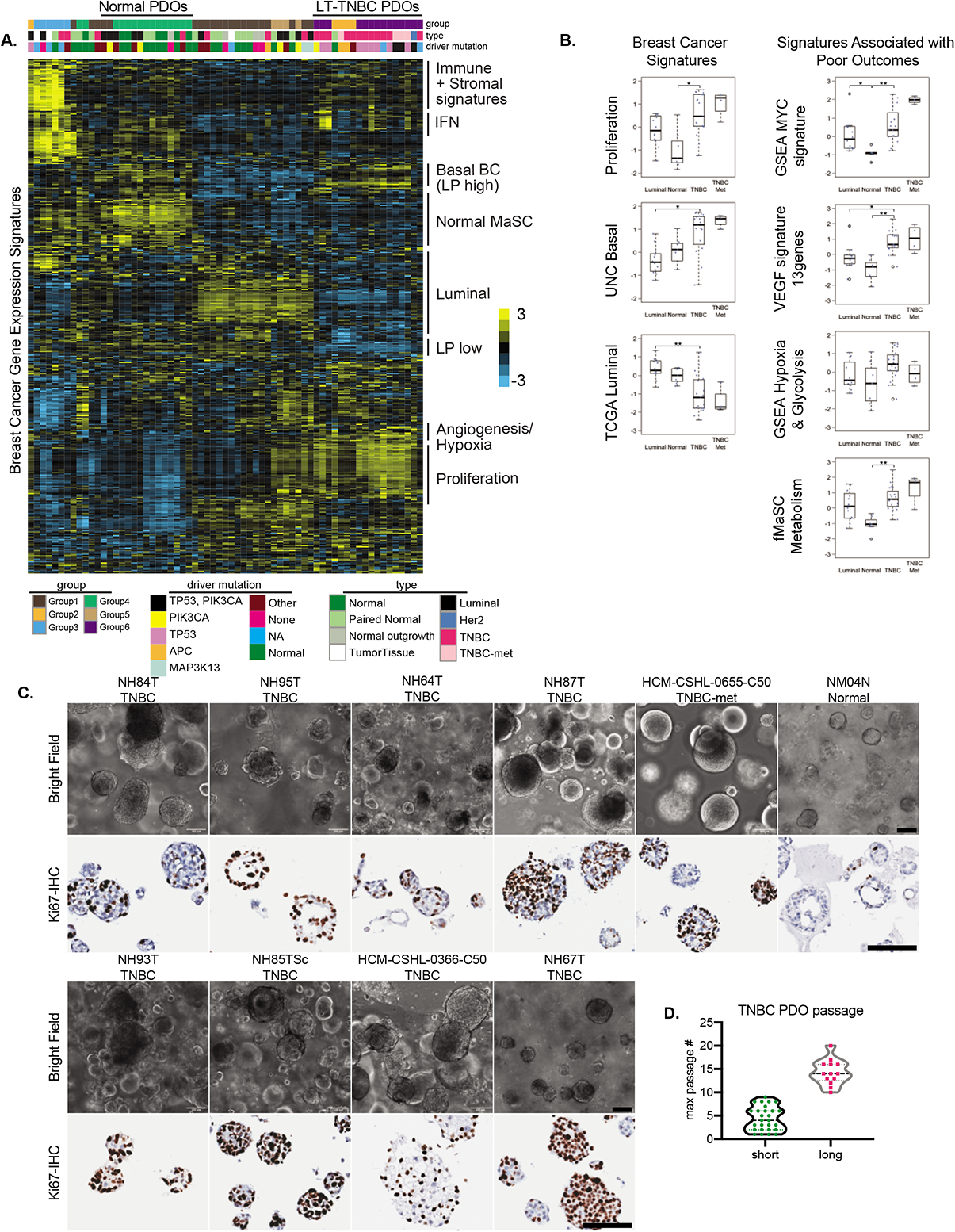
A subset of TNBC organoids recapitulate signatures of aggressive basal-like breast cancers. **A.)** Molecular signatures associated with the different organoid lines. The sample legends are type: Normal= reductive mammoplasty derived normal organoids, Paired Normal= Adjacent or Distal to the tumor paired normal, Normal outgrowth= no pathogenic mutations were found, Luminal= ER/PR+ organoids; driver mutation: Other= trace mutations (see Fig 1E), None= no pathogenic mutations were found, NA= not assessed **B.)** Box-plots showing the breast cancer related and TNBC-specific scores for various gene signatures associated with poor outcomes. Each dot represents a different PDO; Luminal N=12, Normal N= 7, TNBC N= 19, TNBC met= 4. Differences in experimental groups were compared using Kruskal-Wallis test followed by pairwise comparisons using Wilcoxon rank-sum test. Bonferroni-Holm method was used to adjust the family-wise error (** adjusted p-value < 0.005, * adjusted p-value < 0.05) **C.)** Light microscopy images of the various TNBC-and normal (NM04N) derived organoid lines, along with Ki67-IHC, scale bars=100µm **D.)** Distribution of maximum passage numbers tested for the various TNBC PDOs. Long-term cultures (long) are defined by p>10 with continued expansion

Groups 2 and 6 are comprised mostly of TNBC organoids and all organoid lines in these groups could be propagated to long-term cultures (>passage10, labeled LT-TNBCs) and showed continued expansion (Fig 3A, S2A). The only exception is the NH48N normal sample, which was confirmed to be mostly tumor by copy number analysis and identical to its counterpart NH48T (Fig S2B). The most prominent signatures of Groups 2 and 6 correspond to basal-like breast cancer gene-sets that are also defined by luminal progenitor (LP) like signatures (Fig 3A-B, S2C-D). LP-like gene expression has previously been shown to be associated with basal-like breast cancers (40). The organoids in this group also showed enrichment of proliferation signatures (Fig 3A-B) and the majority had a *TP53* mutation which was in conjunction with the *PIK3CA* mutations for three samples (HCM-CSHL-0773-C50, HUB4T and HCM-CSHL-0155-C50) (Fig 3A, S2A). We also observed a MYC amplification signature in this subgroup (Fig 3B) that was accompanied by the copy number amplification of *cMYC* in many of these PDOs (Fig S2E). Additionally, gene sets associated with hypoxia, glycolysis, angiogenesis and fetal MaSC (fMaSC) metabolism signatures were also enriched in LT-TNBCs (Fig 3A-B, S2C). The *VEGF* 13-gene signature showed high correlation with basal-like breast cancers (29), has prognostic significance and is associated with poor outcome in breast and other cancers (41). Similarly, fMaSC metabolism signature is a refined 8-gene signature, which primarily comprises genes associated with glycolysis and fatty acid metabolism, and was previously shown to be associated with TNBCs and metastatic TNBC lesions (42). This suggests that the PDOs with these signatures are likely associated with tumors linked to poor outcomes.

In order to understand the complexity of organoid cultures we profiled the growth properties of various TNBC and normal organoids. TNBC organoids appear as densely filled balls of cells which is in stark contrast to the normal organoids that have a predominant acinar structure with a central lumen and some organoids that were filled (NM04N in Fig 3C). The TNBC-organoids showed varying degrees of propagation in culture where some lines could be propagated for over 10 passages (LT-TNBCs) with continued expansion while many short-term culture lines dropped out of culture at various points (Fig 3D, S3A) and had limited material to assay. Low starting material, limited proliferation, normal and stromal outgrowths were among the primary reasons for short term culture growth (Fig S3A). We focused on the LT-TNBCs for all downstream analyses. IHC for the proliferation marker, Ki-67, in confluent PDO cultures showed large variability of expression amongst the various organoid lines and also within each culture (Fig 3C-lower panels). In concordance with expression signatures (Fig 3B), LT-TNBC organoids were highly proliferative compared to normal and luminal organoids (Fig 3C, S3B-C). There was a lot of heterogeneity within organoid cultures with some organoids being mostly comprised of proliferating cells, for instance NH85TSc and NH95T, while other cultures were more mixed (Fig S3C). Normal-derived organoids typically had <10% of proliferating cells per organoid and showed luminal-basal organization with a hollow lumen, while TNBC-derived organoids tend to be highly proliferative and undifferentiated with little to no observed cellular organization (Fig S3C). When seeded as single cells, we observed a range of organoid formation amongst the various lines (Fig S3D) which did not always correlate with the proliferative index of the PDOs, and might be due to the requirement of cell-cell contact in some but not all PDOs that is apparent when seeding at low density as single cells.

Of note, none of the TNBCs propagated in culture showed expression profiles of the more mesenchymal-like claudin-low subgroup (6), possibly because the culture conditions are more favorable towards the propagation of epithelial cells. Overall, the gene expression signatures of the normal and tumor-derived organoids depict the expression profiles of luminal and basal-like breast cancers. The TNBC organoids that can be propagated to long-term cultures (LT-TNBCs), Groups 2 and 6, had very classic basal-like breast cancer signatures and were associated with proliferation (Fig 3A-D), hypoxia (Fig 3A-B) and *c-MYC* amplification gene expression signatures (Fig 3B, S2E). Furthermore, about 40% of basal-breast cancers exhibit *c-MYC* amplification (26) and the majority of our LT-TNBC organoids show an enrichment of *c-MYC* (Fig S2E) signature suggesting that our organoid system results in long-term expansion of TNBCs that are *c-MYC* driven and have a LP-like basal breast cancer signature.

As a proof-of-concept, we tested whether these PDOs were amenable to drug response assays, and performed dose response curves using 6 different PDO lines with three different drugs (Fig S3E). TNBC PDOs (Fig S3D) showed high sensitivity to the cytotoxic chemotherapeutic agent paclitaxel compared to the luminal line NH53T and the TNBC-met HCM-CSHL-0773-C50 (Fig S3E). Conversely, capivasertib (AZD5363), a pan-AKT inhibitor used in PIK3C mutated cancers (43), showed greater sensitivity against the PIK3CA mutated luminal NH53T PDO (Fig S3E). Similarly, afatinib a targeted inhibitor for EGFR, showed the most activity in the EGFR amplified NH84T PDO line (Fig S3E).

### TNBC organoids can recapitulate tumor morphology *in-vivo*

Next, we examined the *in-vivo* tumor forming abilities of these patient-derived cultures. We transplanted organoids from eight different TNBC (LT-TNBCs) lines into the mammary-fat pads of NOD-SCID mice and assayed their tumor formation and metastatic potential over time (Fig 4A). We found striking differences in the tumor forming ability between the different organoid lines (Fig 4B) despite having highly aberrant genomic profiles (Fig 2, 4C). TNBC organoids HCM-CSHL-0366-C50, NH85TSc and NH87T resulted in palpable masses around 30 days and steady tumor formation in 100% of injection sites (Fig 4B, Table S6). NH85TSc-derived tumors grew rapidly and resulted in visibly necrotic masses that needed to be resected at ∼100 days. HCM-CSHL-0366-C50 resulted in primary tumors that also developed fluid filled cysts on top of the tumors that drained on their own. NH87T and HCM-CSHL-0655-C50 are the AdCC-like primary tumor and lymph-node metastasis from the same patient. Interestingly, while NH87T formed tumors rapidly, HCM-CSHL-0655-C50 did not and the tumors remained small (Fig 4B, Table S6), despite both of these lines having equivalent growth properties *in vitro* (Fig 3, S3). This observation was consistent for the two independent transplant experiments done with different passage organoids (Table S6). NH95T resulted in small tumor masses at multiple injection sites that were slow to grow while NH84T and NH93T had small tumors observable only in fat-pad histology sections of some sites and NH64T did not result in any primary tumors (Figs 4B, S4A, Table S6) despite being highly genomically aberrant (Figs 4C, 1-2) and having high proliferation and organoid formation rates *in-vitro* (Figs S3C-D).

**Figure 4.**
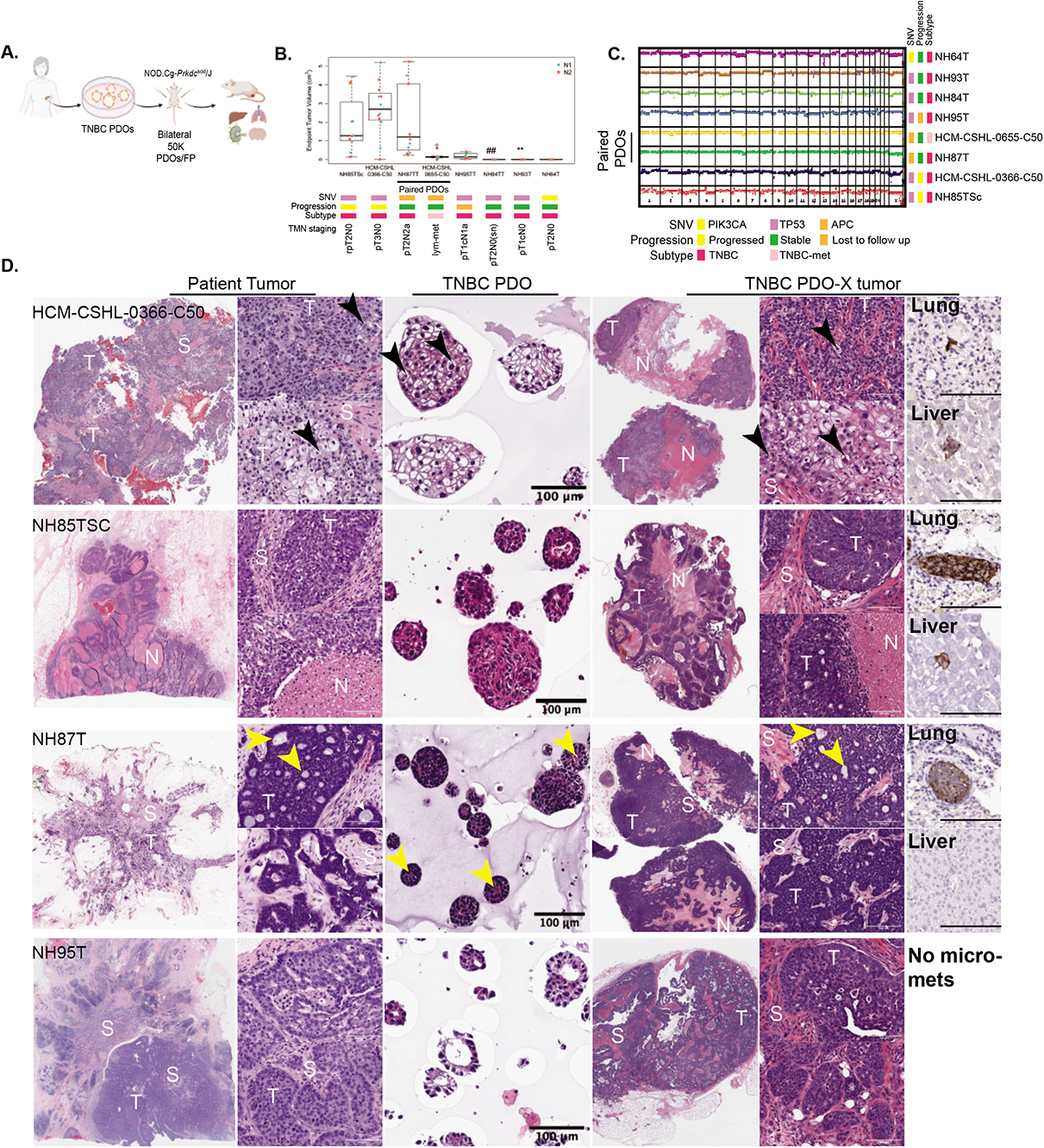
TNBC Organoids can recapitulate tumor morphology and progression *in-vivo.* **A.)** Overview of PDO xenotransplant experiment using TNBC PDOs. **B.)** Box plots showing the end point tumor volume for the various organoid lines transplanted into the fat-pads of NOD-SCID mice. Each dot represents tumor volume from one injection, N1= experiment 1, N2= experiment 2. ## For NH84T microscopic primary tumors observed in histology sections from 1/10 sites. ** For NH93T microscopic primary tumors observed in histology sections from 6/10 sites. TMN staging: pathologic TMN staging from patient pathology report (Table S1) **C.)** CNV profiles of the PDOs lines selected for *in vivo* transplant experiments **D.)** H&E images of the paired patient tumor tissue, TNBC patient-derived organoids (PDO) and xenografts generated form patient-derived organoids (PDO-X). S= Stroma, T= tumor, N=necrosis. Black arrows point to the cells with spinous connections with adjacent tumor cells. Yellow arrows point to the pseudo-lumen observed within AdCC-like breast cancers. Last column shows human mitochondria IHC in the lung and the liver from a representative mouse injected with the respective PDO (scale bar=100µm).

Remarkably, the PDO-derived xenograft (PDO-X) tumors resulting from HCM-CSHL-0366-C50, NH85TSc, NH87T and NH95T organoids recapitulated the morphology of the patient tumor despite previously being in culture for up to 18 passages. The tumor and the organoid-xenograft from patient HCM-CSHL-0366-C50 were the most distinct and showed squamous differentiation with pleomorphic cells that were variable in size and had cells with either oval or pale nuclei with prominent nucleoli (Fig 4D). We also observed cells that had thin spinous connections with adjacent tumor cells which were also present in the organoids (black arrows Fig 4D). NH85TSc tumor and organoid-xenografts both showed highly undifferentiated morphology with large necrotic areas and stromal compartments infiltrating in between the tumors (Fig 4D). Lastly, NH87T tumor was a TNBC type with AdCC like features that are characterized by cribriform architectural patterns and pseudo-lumens (31, 44). While the organoid-derived xenograft had a lower stromal composition compared to the patient tumor it still recapitulated the key features of the original tumor, including the presence of cuboidal cells, cribriform architecture and pseudo-lumens (yellow arrows Fig 4D). This particular subtype is also characterized by the expression of CD117 and CK5/6 (45). We performed IHC for CD117 and CK5/6 on PDO-X tumor sections from NH87T and observed positive membrane labeling for CD117 along with areas of high and low CK5/6 labeling as observed in the clinical IHC labeling of the patient slide (Fig S4C).

While NH95T formed tumors at a much lower efficiency (Fig 4A, B), the tumors that did form had morphological similarities to the original patient tumor (Fig 4D). The patient tumor cells showed an organization pattern which was recapitulated in the organoid-derived xenografts and to some extent in the NH95T organoids (Fig 4D). Of note, patients from whom the fast-growing organoids were derived, i.e., HCM-CSHL-0366-C50, NH85 and NH87, presented with poor diagnoses and outcomes (TMN staging in Fig 4B). Patient HCM-CSHL-0366-C50 had rapid metastasis to the brain and succumbed from the disease, patient NH85 had local recurrence in the lymph node and patient NH87 presented with a positive lymph node at initial diagnosis (Table S1). Using IHC with a human mitochondrial antibody we only observed micro-metastasis and single-cell metastasis in our experiments (Fig 4D, S4B, Table S6), however, altering the experimental conditions such as: prolonging the end-point, using a more immunocompromised NOD/SCID gamma (NSG) mouse model or tail-vein injections might result in more metastatic lesions and will be examined in future studies. Thus far, our data shows that TNBC PDOs recapitulate the tumor intrinsic properties of the original tumors at genomic, transcriptomic and morphological levels.

### TNBC derived organoids are enriched for luminal-progenitor-like cells

To fully assess the utility of PDOs as cancer models, we next asked what were the cell types represented within these organoids and how did they differ from the normal derived PDOs. In order to profile the cell-types present within the normal and TNBC organoids, we used a combination of flow cytometry and single cell RNA-seq (scRNA-seq). We assayed for mammary epithelial lineages using EPCAM and CD49f as luminal and basal cell markers respectively (21,22,40,46). As previously shown (22), the normal derived organoids recapitulate the EPCAM+ luminal lineage, EPCAM+/CD49f+ luminal progenitor cells and EPCAM-/CD49f+ basal cell lineages (NM07NL in Fig 5A). The populations slightly fluctuated between the different normal lines and passages but overall, the different lineages were observed in all normal organoids (Fig 5A-C, S5A, Table S7). TNBC organoids were less heterogeneous in the expression of these markers, and as observed by RNA-seq signatures (Fig 3B, S2D), showed an enrichment of luminal progenitor-like cells (Fig 5A-B) which had co-expression of EPCAM and CD49f but showed slightly different and patient-specific expression patterns of these markers (Fig 5A, S5A). TNBC organoids also showed a higher gene expression of EPCAM and CD49f (gene *ITGA6*) compared to the normal organoids (Fig S5B). Some TNBC PDOs, for example NH84T, had a more luminal cell flow-cytometry profile, despite clustering with the other LT-TNBCs at the transcriptome level. Interestingly, metastatic TNBC lines HCM-CSHL-0773-C50 and HUB4T also had an enrichment of more luminal-like cells (Fig 5B, S5D). NH85TSc which was derived from a patient that later relapsed and HCM-CSHL-0366-C50, which showed rapid progression in the patient, also had a more luminal cell profile with higher percentage of EPCAM only cells (Fig 5A-B) despite showing gene signatures of the basal-like BCs. The EPCAM/CD49f profile of the individual lines was stable overtime at different passages (Fig S5C). Additionally, while always observed in the normal-organoids, we rarely observed CD49f only cells in the TNBC-organoids (Fig 5A-B, S5A, D). We performed longitudinal analysis for one of the patient-derived organoid lines and observed an enrichment of the tumor LPs in culture over time in early passages (Fig 5C) which coincided with a more pronounced copy number profile of the TNBC organoids (Fig 5D) suggesting LP-like cells being the predominant cancer cell population in TNBCs and potentially the cell of origin of these cancers.

**Figure 5.**
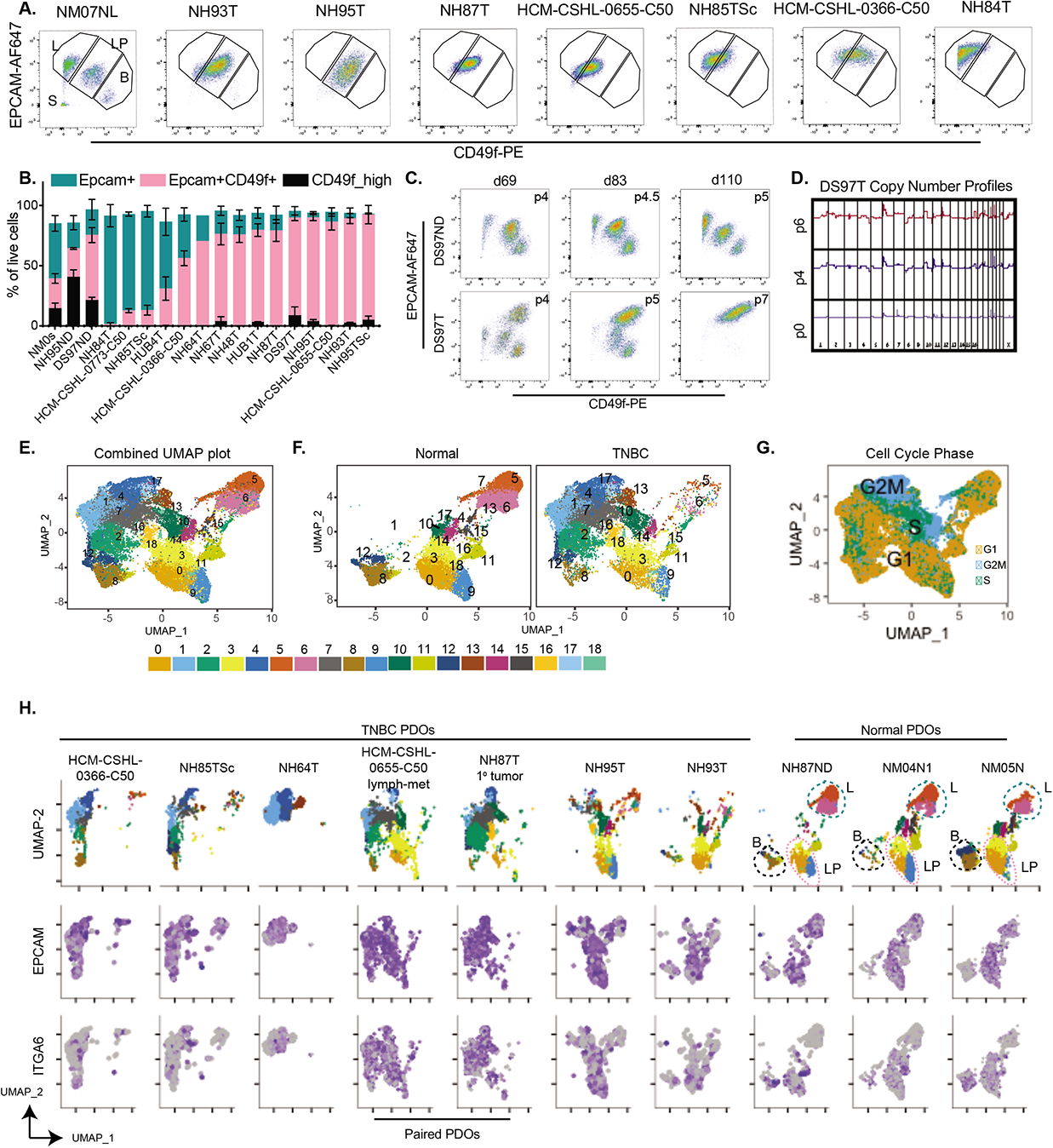
TNBC organoids are enriched in luminal progenitor-like cells. **A.)** Representative flow-cytometry plots for normal derived organoid (NM07NL) and various TNBC organoids stained for CD49f-PE on the x-axis and EPCAM-AF647 on the y-axis. The gates are subsets of live single cells within each organoid line and represent various cell types of the mammary epithelium. L=EPCAM-high mature luminal cells, LP= EPCAM+CD49f+ luminal progenitors, B= CD49f+ basal cells, S= stromal compartment. **B.)** Quantitation of the L, LP and B gates in panel B for multiple TNBC and normal organoid lines over multiple passages (see Table S7). Data-points are plotted as mean **±**SEM using GraphPad Prism. NM0s: comprises multiple normal mammoplasty derived organoids from different patients **C.)** Flux of the epithelial cells during the early passages of organoid derivation for normal distal (DS97ND) and TNBC tumor (DS97T) samples from the same patient **D.)** Copy number plots of the TNBC organoids DS97T over multiple passages **E.)** UMAP plot of batch corrected scRNA-seq data from 3 normal and 7 TNBC organoids. Numbers on the plot represent cluster IDs. **F. & G.)** UMAP plot in A but **F.)** separated by the normal and tumor samples and **G.**) colored by cell cycle. **H.)** UMAP plot in E split by individual tumor and normal samples and showing normalized expression of *EPCAM* and *ITGA6* (gene encoding CD49f) expression patterns in individual cells.

In order to better understand the cellular composition of these organoids, we performed scRNA-seq analysis on three normal and seven TNBC organoid lines. The study was done in three experimental batches (Table S7) and processed where each of the samples was quality controlled and filtered to remove cells with high mitochondrial gene content and low gene identification (i.e. dead cells). The filtered matrix was SC-transformed using Seurat (47) and the samples were integrated to account for the different batches (Fig 5E). Clustering of data identified 19 clusters (Fig 5E) with some clusters being highly representative of the normal organoids (clusters 0, 3, 5, 6, 9, 11 in Fig 5F) while other clusters were largely comprised of tumor cells. Tumor organoids were distinct from normal, with some overlap in clusters 0, 3 and 9 (Fig 5F-H). Expression profiles of *EPCAM* and *ITGA6* (gene encoding CD49f) (Fig 5H) recapitulated the data from flow cytometry (Fig 5A-B) where tumor organoids are predominantly of the luminal progenitor nature as measured by co-expression of *EPCAM* and *ITGA6* (CD49f) while the normal organoids recapitulate the three broad mammary epithelial lineages: mature luminal (EPCAM+), luminal progenitors (LPs, EPCAM+/CD49f+) and myoepithelial/basal-like cells (Epcam-low/CD49f+). Unlike the normal organoids, all tumor cells were *EPCAM*+ and a subset of those had low *ITGA6* (Fig 5H). In concordance with the flow cytometry data (Fig 5A), the *ITGA6*-low cells were largely present in HCM-CSHL-0366-C50 and NH85TSc. NH93T and the NH95T organoid line had similar profiles and occupied similar space as the normal LP cells (Fig 5H). Lines NH87T and HCM-CSHL-0655-C50 are paired primary and lymph node metastasis samples from the same patient and occupied very similar spaces with some overlap with normal LP cells (Fig 5H), while NH64T, NH85TSc and HCM-CSHL-0366-C50 are very distinct from the normal lines. Interestingly, there was a significant overlap between the clusters identified between NH85TSc and HCM-CSHL-0366-C50 samples (Fig 5H) in keeping with their transcriptome similarity (Fig 3A, S2A) and rapid tumor progression *in vivo* (Fig 4A).

This data builds on previous studies that have correlated a LP-like expression signature with basal-like breast cancers (40) and have shown that LP cells are the cell of origin of BRCA-mutated basal breast cancers (48). We suggest that LP-like cells are the possible cell of origin for a broader subset of TNBCs. Interestingly, a higher percent of EPCAM+ only cells seems to be associated with a greater degree of disease progression, however, this needs to be further investigated.

### Tumor LP-like cells exhibit altered expression and have an upregulation of NOTCH and MYC downstream pathways

Since TNBC organoids had a large proportion of LP-like cells that seemed distinct from the normal LP cells (Figs 5, S5) we performed integrated single cell analysis by SC-transforming individual samples and performing an anchor-dependent integration for all individual samples. We identified thirteen clusters between the tumor and normal cells (Fig 6A) out of which clusters 6, 8, 10 and 9 represented cell cycle clusters (Fig S6A) with cluster 6 representing a population of G1-S phase cells, cluster 8 representing S-phase cells, cluster 9 S-G2M transition cells and cluster 10 marking G2M cells while the remaining clusters represented G1 cells (Fig S6A-C). The cell clusters identified represented biologically meaningful cell-types recently annotated by single cell sequencing of adult human breast epithelium (49). Cell types identified included: XBP1, AGR2 expressing mature luminal cells (Mat lum, cluster 1), APOE, KRT6A expressing basal stem-like cells (basal SC, cluster 4), TAGLN, TIMP1 expressing myoepithelial cells (Myo, cluster 3), SLPI expressing luminal progenitor cells (LP, cluster 7) and LTF expressing secretory cells (LP sec, cluster 2) (Fig 6B, S6C) (49). To further validate the lineage specification in organoids we overlaid the published mammary epithelial gene-sets (40) and computed a score for each of the three mammary epithelial cells. The normal cell clusters showed a much stronger cell type enrichment score while the tumor cells despite being predominantly EPCAM+/CD49f+ showed a more diffused enrichment of these signatures, suggesting some cell type heterogeneity within these tumor organoids (Fig 6C). In normal organoids, Clusters 2,7 and 11 had a higher enrichment for luminal progenitor (LP) cell score, while cluster 1 showed an enrichment for mature luminal (mature Lum) cells and cluster 3 was predominantly of the basal mammary stem cell (Basal SC) compartment (Fig 6C, S6C). The tumor organoids showed varied expression profiles of various luminal/basal markers (Fig S6D) and a diffused cluster specific enrichment of lineage scores (Fig 6C, S6E), suggesting that while they have lost proper cell type specification, the clusters identified using this integration approach have some similarity to the normal cell lineages.

**Figure 6.**
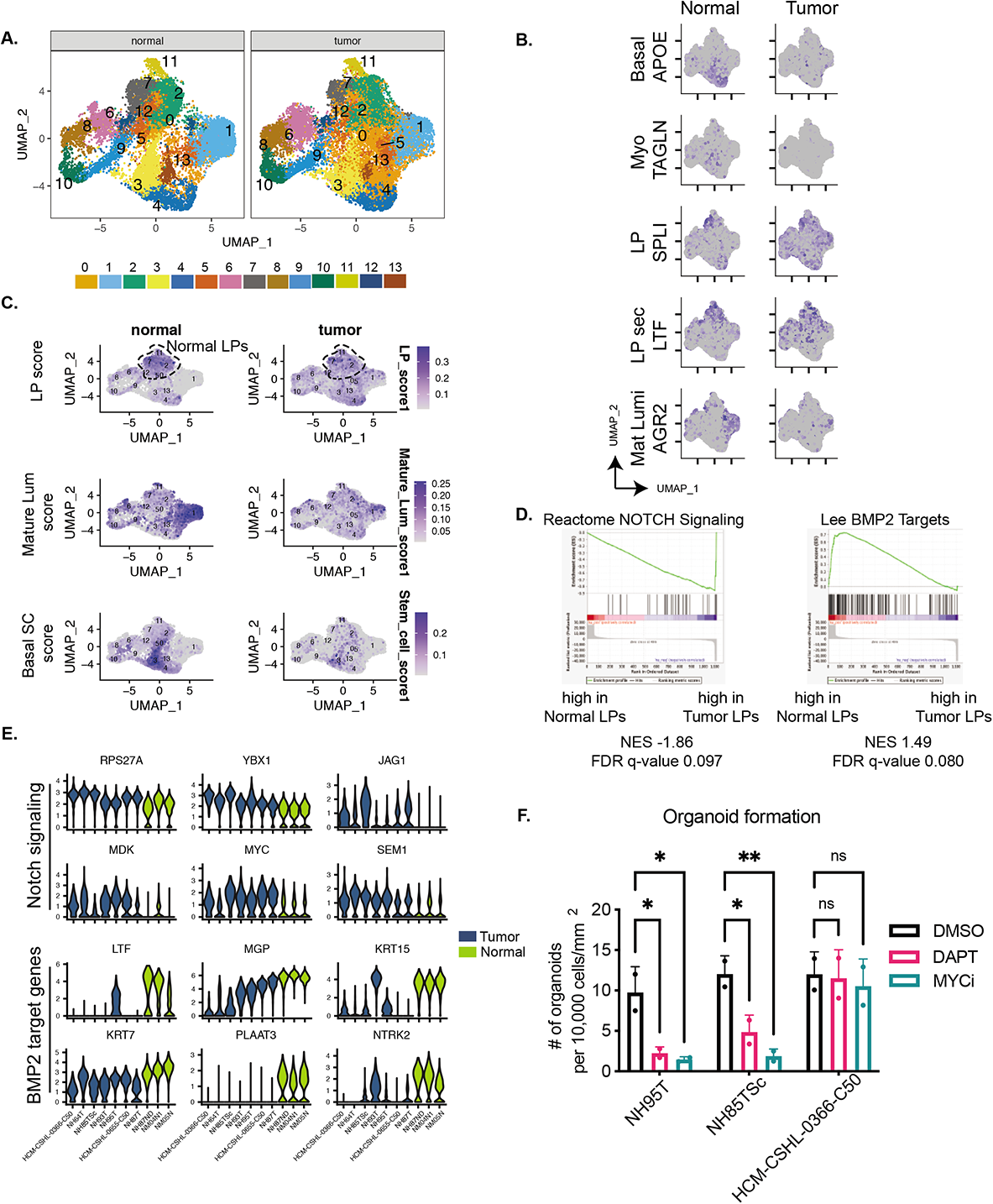
Tumor LP-like cells exhibit altered gene expression and have an upregulation of NOTCH and MYC downstream pathways. **A.)** UMAP plots for Integrated scRNA-seq data for all samples. The numbers indicate cluster IDs. **B.)** Marker expression of various cell type specific genes in the adult human breast epithelium (49). **C.)** Plots showing combined scores for the three mammary epithelial lineages: LP score: Luminal Progenitor score, Mature Lum score: Mature luminal score, MaSC score: Mammary stem cell score (40). Dashed region indicates LP clusters 2,7 and 11 that were used to perform differential expression analysis between normal and tumor LPs. **D.)** GSEA plots showing enrichment of the differentially expressed genes between normal and tumor LPs. **E.)** Violin plots showing combined expression in clusters 2,7 and 11 of the leading-edge NOTCH signaling genes and BMP2 target genes as identified in E. **F.)** Organoid formation from single cells. Significance was assessed by two-way ANOVA ns= not significant, ** pvalue<0.005, * pvalue<0.05

With the aim to identify mechanisms that underlie the tumor luminal progenitor-like (LP-like) cells in TNBCs, we performed a differential expression analysis between tumor and normal cells of clusters 2, 7 and 11. We identified 1103 significantly differentially expressed genes (p_val_adj<0.05) between the tumor LPs vs normal LPs (Table S8). GSEA on the differentially expressed genes showed an enrichment of NOTCH signaling related genes in the tumor LPs versus an enrichment of BMP2 targets in the normal LPs (Fig 6D). Leading edge genes from these gene sets showed a consistent downregulation of NOTCH signaling related genes, *RPS27A*, *YBX1*, *JAG1*, *MDK*, *MYC* and *SEM1*, and an upregulation of BMP2 target genes, *LTF*, *MGP*, *KRT16*, *KRT7*, *PLAAT3* and *NTRK2* in normal LPs from all three patients (Fig 6E). MDK, JAG1, YBX1 are ligands of NOTCH1, while *MYC* and *RSP27A* are downstream targets. While NOTCH activity was present in normal LP cells, it was more pronounced in the TNBC organoids (Fig S6F).

A further investigation into differentially regulated pathways showed upregulation of genes involved in ribosomal biogenesis and translation in the tumor LPs. We performed motif enrichment analysis on the differentially expressed LP genes and found a significant enrichment of MYC binding sites in the tumor LP expressed genes (Fig S6G). We used published MYC signatures to assess MYC activity in tumor and normal organoids and found a higher ubiquitous enrichment of MYC activation in tumor organoid cells, while in normal organoids it seemed to be higher in the proliferating cell clusters (Fig S6F). Similarly, while the NOTCH pathway was hyper-active in tumor LPs it was ubiquitously active across all tumor cells (Fig S6F). We tested whether NOTCH and MYC activation was required for organoid formation and seeded NH95T, NH85TSc and HCM-CSHL-0366-C50 as single cells in regular organoid growth medium in the presence of DAPT (NOTCH inhibitor) or MYCi975 (MYC inhibitor) (50) and allowed for organoids to form for 12 days. We saw a significant reduction in organoid formation in the presence of both MYC and NOTCH inhibitors for NH85TSc and NH95T but not for HCM-CSHL-0366-C50 (Fig 6F). While HCM-CSHL-0366-C50 did not show reduction in the number of organoids formed, we did observe a significant difference in organoid size with DAPT and MYCi975 suggesting growth defects on inhibition of these pathways in all lines (Fig S6H). We further tested whether inhibition of NOTCH and MYC is necessary for the formation of normal PDOs. Since, normal PDOs have multiple populations we flow-sorted the basal stem-like CD49f+ve population and EPCAM+/CD49f+ LP cells and seeded them as single cells in the presence of DAPT or MYCi975 (Fig S6I). While DAPT had no effect on generation of normal PDOs from both LP and basal stem-like cells, MYC inhibition resulted in no organoid formation.

Thus, our data shows that TNBC organoids have an underlying LP expression signature which is driven by LP-like cells that exhibit altered expression from normal LPs by hyperactivation of NOTCH and MYC signaling. These pathways are necessary for the formation of some, but not all, TNBC organoids and when perturbed result in proliferation defects in all tumor lines examined.

### TNBC organoids are comprised of heterogenous cancer cell populations

Since normal organoids have clearly defined lineages, the clustering of the integrated dataset (Fig 6) was largely driven by the normal organoids. To assess the heterogeneity that exists within TNBC organoid cultures we performed a similar integrated analysis on the TNBC organoids only and identified 13 different cell clusters (Fig 7A, S7A). Clusters 3,9,6,8 and 5 correspond to the different cell cycle phases (Fig 7A) and the remaining clusters are G1 cells (Fig 7A-B). We identified the markers that uniquely define each of the clusters (Fig 7C, Table S9) and performed GSEA on the marker genes to identify the phenotypes associated with each cell cluster (Fig 7D). Cluster 0 represented a mixed cell cluster with some enrichment of mature luminal-like cells. Cluster 1 was defined as Basal-like 1 as it showed a positive enrichment of a basal breast cancer gene-set including specific markers *SAT1*, *GABRP*, *TM4SF1*, *TTYH1*, *KRT16* and *KRT6A* genes (Table S9). Cluster 2 is mesenchymal-like due to the selective expression of mesenchyme genes including *CCN2*, *TPM1* and *LAMB1*. Cluster 7, MGP-high cluster, is a relatively distinct cluster with a very specific high expression of *MGP* (Matrix Gla Protein) in all TNBC organoid lines (Fig 7C, S7E). MGP is normally expressed in smooth muscle cells perhaps suggesting that these cells might have contractile abilities. Cluster 12 had a very high expression of ribosomal and translation related genes (Fig 7C-D). Clusters 4,10 and 11 are the most distinct of the tumor cell clusters and are identified by the unique expression of certain genes. Cluster 10 represents an NFKB-active cluster that was also found in normal organoids (Fig S6C), albeit with some differences in gene expression. As in normal organoids (Fig 6A, S6B, 8C). this cluster shows the expression of *CXCL1*, *CXCL3*, *NFKBIA* etc. Cluster 11 is also defined by a basal breast cancer signature (Fig 7C-D), is labeled Basal-like 2 and shows a selective expression of *KRT17*. Cluster 4 is a hypoxia cell cluster that shows selective high expression of *EGLN3*, *NDRG1*, *VEGFA* and likely represents cells at the center of these organoids (Fig 7C-D). This cluster is represented by a higher hypoxia score (Fig 7C-D), which interestingly correlates with basal stem cell (Basal SC) signatures and shows comparatively low activity of MYC and NOTCH pathways (Fig 7E and S7D-E). We also noted that the cluster specific genes for all clusters were fairly well conserved between the different PDO lines (Fig S7F). Overall, this data suggests that while the TNBC organoids retain patient intrinsic properties (Fig 1-4) there are common cell signatures that define the cell types within these organoids.

**Figure 7.**
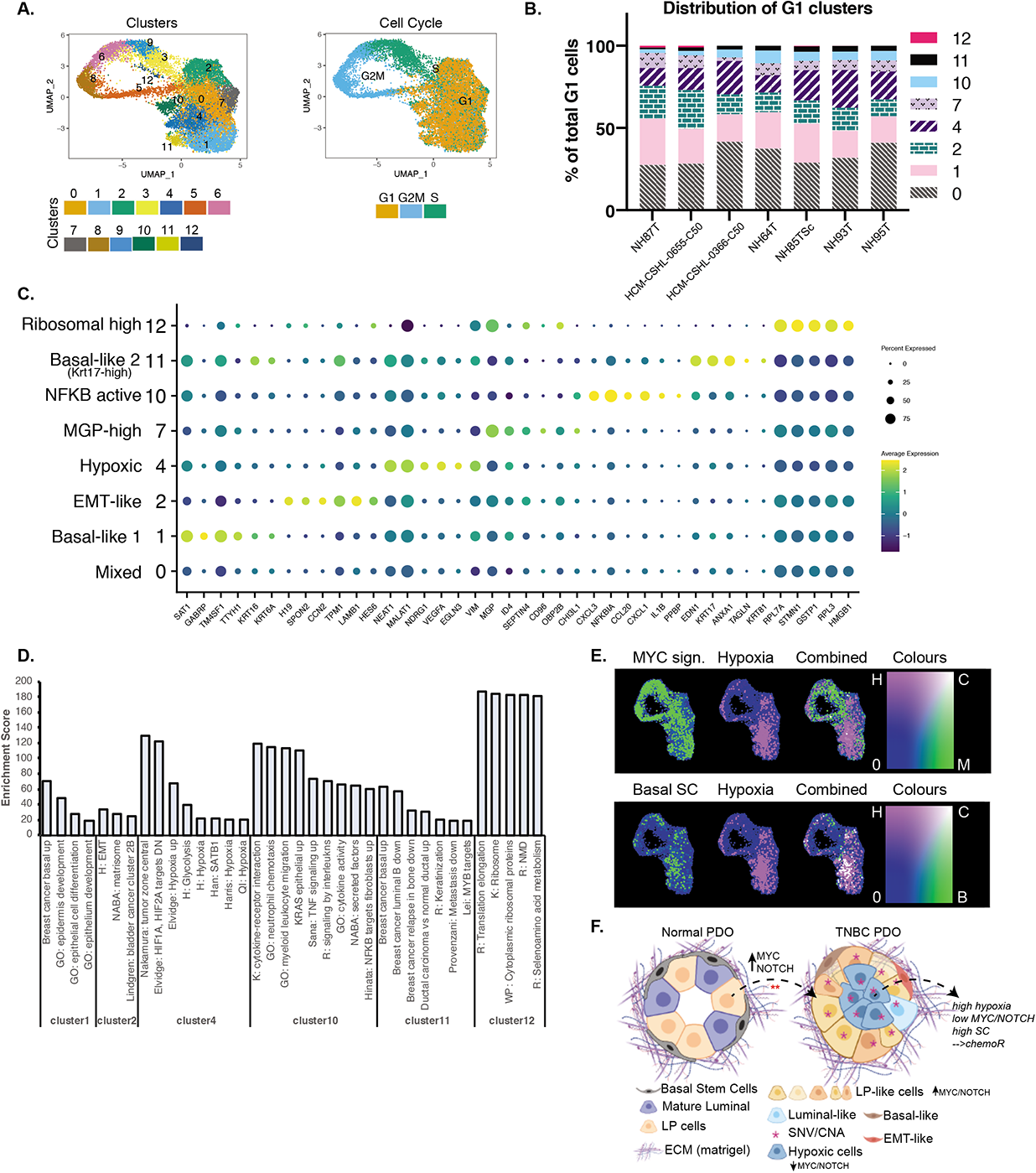
TNBC Organoids are comprised of heterogenous cell populations. **A.)** UMAP plot of TNBC only integrated scRNA-seq data showing clusters identified and cell cycle phases. **B.)** Distribution of cells in each of the G1 clusters identified per organoid line. **C.)** Dot-plot showing the marker genes for each of the G1 clusters and the associated phenotypic identity of that cell cluster. **D.)** Enrichment scores from GSEA of each of the G1 clusters that showed strong enrichment of some specific pathways or phenotypes. Enrichment score is represented by -10*NES*padj.value. **E**.) Combined gene set scores for the various phenotypes. Top panel: green= MYC signature, pink= Hypoxia signature. Bottom panel: green= Basal mammary stem cell (SC) signature, pink= Hypoxia signature. White represents positive correlation of the two signatures. **F.)** Schematic (created using BioRender.com) showing the cellular composition and heterogeneity observed in normal vs TNBC PDOs when cultured in matrigel. TNBC PDOs retain the tumor SNV/CNA profiles, are largely comprised of LP-like cells that might have originated from normal LP cells by the hyperactivation of NOTCH/MYC pathways. TNBC PDOs also have cells with signatures of hypoxia which is anti-correlated with NOTCH/MYC and positively associated with basal mammary stem cell signatures.

The distinct gene expression patterns within these cell clusters also suggests that the TNBC organoids are comprised of multiple cell types that are known to be involved in tumorigenesis, tumor progression and metastasis, and can therefore be used as models to study the various aspects of breast cancer progression.

## Discussion

We developed a patient-derived biobank of normal and breast cancer organoids from a diverse group of patients with a focus on the highly aggressive TNBC subtype. Our biobank is heterogenous in terms of ethnic/racial backgrounds, patient age and breast cancer subtypes. A comprehensive genomic, transcriptomic, and cellular characterization of these PDO models demonstrate their faithful recapitulation of the patient tumor intrinsic properties and hence validates them as cancer models to study various aspects of breast cancer progression and treatment.

Patient-derived organoids are an exciting step towards personalized medicine with a promise to be used for real-time drug screens for guiding patient treatment (14). When successfully established, breast cancer PDOs retain the genomic and transcriptomic features of breast cancers and their parent tissues. We found that, the TNBC organoids that showed long-term robust growth had activated MYC signaling, an LP-like gene expression signature and were overwhelmingly composed of LP-like cells. As these are among the most aggressive forms of breast cancer, organoid models appear to be an exciting avenue for their study. In addition to replicating the tumor specific features *in-vitro*, TNBC PDOs when transplanted into NOD/SCID mice generated tumors with remarkable morphological similarity to that of the original patient tumors, despite being in long-term organoid culture for up-to 18 passages. Long-term cultured organoids, thus, maintain the intrinsic ability to represent the tumor from which they were derived when placed in an *in vivo* environment. Interestingly, in our study not all PDOs readily generated primary tumors *in vivo* despite being highly proliferative and genomically aberrant organoids. The association of PDOs that generated tumors rapidly, with worse outcomes and diagnosis, suggests that there might be a biologically relevant explanation for this observed difference and must be investigated further along with long-term patient follow up information. Patient-derived xenograft (PDX) models also retain the histopathology of the original patient tumors (51, 52) and a recent pre-print study showed successful derivation of breast cancer organoids from PDX models (53). Our models are complementary to this system and represent an opportunity to derive PDOs first, perform *in-vitro* assays and drug screens, followed by PDOX derivation for *in-vivo* validation studies.

While successfully established PDOs faithfully recapitulate the patient tumor properties, the efficiency of establishment of these cultures is currently less than ideal. The common challenges we faced during establishing organoid cultures included normal outgrowth in some tumor organoid cultures, stromal outgrowth, non-proliferating or dormant tumor cells, or limited growth potential *ex-vivo*. Since the medium composition for tumor organoid growth also supports the culture of normal mammary organoids (22) the purity of the starting tumor tissue might govern the time to generation of tumor-only organoid cultures and early passages should be meticulously tested at a genomic level before any pharmacological studies are carried out with these models. Furthermore, some subtypes of breast cancer might be challenging to culture including the less proliferative Luminal A subtype or the more mesenchymal claudin-low group of TNBCs. Efforts to culture tumor-organoids in reduced complexity medium (53), might alleviate some of these challenges and needs to be examined in future studies. The PDO culture conditions are comprised of complex growth factors including, EGF, FGF10, Neuregulin, R-spondin etc. (20), and it remains to be tested whether the removal or alteration of these components would be beneficial in selecting out cancer cells over normal epithelial cells, resulting in higher efficiency in the establishment of TNBC PDOs. However, the simultaneous growth of normal and TNBC organoids in the same culture conditions allows for comparison of similar cell types within these organoids.

We used an integrated scRNA-seq approach to compare normal LP cells with the tumor LP-like cells and identified hyperactivation of NOTCH and MYC signaling in the tumor compared to normal LPs. LP cells were previously shown to be the cell of origin of BRCA1 mutated basal-like BCs (40, 48) and are speculated to be involved in tumorigenesis of all basal-like BCs due to similarity of gene expression profiles (40). Our data provides strong support for this hypothesis by showing that TNBC PDOs are largely comprised of LP-like cells and suggests that hyperactivation of NOTCH and MYC signaling might be relevant in tumorigenesis from LP-cells (Fig 7F). Mouse studies have shown that Notch signaling is important in the maintenance of luminal lineage of the normal mammary gland (54–57) and overactivation of Notch in luminal progenitors resulted in hyperplasia and acquisition of self-renewal properties (54). Notch activation drives the luminal fate specificity in normal mammary gland and can reprogram committed basal mammary cells to an ER-luminal cell fate via multipotent embryonic cell states (55, 58), suggesting a role for Notch in fate specification and cellular plasticity. While our data provides strong support for hyperactivation of NOTCH signaling in LP-like cells in TNBCs it remains to be tested whether these cells arose from activation of NOTCH in normal LP cells or from reprograming of normal basal cells. Our data suggest that activation of MYC along with NOTCH might also have a role in this transformation. Given the complementarity of the NOTCH and MYC pathways, further studies are needed to gain better resolution of this mechanism. The PDO system, thus, can be exploited for identification of dysregulated pathways in cancers which can then be perturbed in the normal PDOs to understand aspects of the origins of cancers.

A detailed analysis of the cell type heterogeneity of TNBC organoids also revealed the presence of MYC/NOTCH-activity low and hypoxia high cells amongst various other cancer relevant cell types. Recent studies have shown that cancer cells can enter a MYC-low diapause-like state of dormancy upon chemotherapy treatment and result in therapy escape (59, 60). Furthermore, single cell analysis from TNBC tumors with residual disease after chemotherapy treatment showed an enrichment of hypoxia, angiogenesis, EMT and ECM degradation related genes in the persistent tumors (61). We hypothesize that in PDOs with these expression signatures, the cells that exhibit a MYC-low/hypoxia-high profile are the likely candidates for chemotherapy escape (Fig 7F). Additionally, fate mapping and other studies have shown that tumor cells exposed to hypoxia have a higher tendency to result in metastasis (62, 63). It remains to be tested to what extent these MYC-low/hypoxia-high cells contribute towards metastasis and/or chemotherapy resistance, and these patient-derived organoids will be an invaluable tool to answer these questions.

In summary, we developed a diverse biobank of BC organoids with a focus on TNBC-derived organoids. We have thoroughly characterized these models as valid systems that mimic the various aspects of patients’ tumors, including genomic alternations, transcriptomic signatures, cell type specificity and morphological characteristics. Comparison of TNBC and normal-derived organoids provides important insights into the mechanisms regulating tumorigenesis that can then be validated by perturbation in normal PDOs (21). This comparison can also be used to identify novel tumor specific targets that may play an important role in tumor growth and progression (64). These next generation cancer models and the data derived from them offer vast utilities and can be used for drug-screens, co-culture experiments, metabolomics, and fate mapping studies to better understand the mechanisms driving cancers and for identifying more specific treatment options.

## Acknowledgements

We would like to thank members of the Spector lab for critical discussions and advice throughout the course of this study. We thank Bodu Liu for critical review of this manuscript. We also thank the various members of NCI-Human Cancer Model Initiative (HCMI) for general advice and inputs on organoid cultures. We acknowledge the CSHL Cancer Center Shared Resources (Animal, Flow Cytometry, Histology, Microscopy, Organoid, Next-Gen Sequencing, and Single Cell Biology) for services and technical expertise (NCI 2P3OCA45508). The sequencing analysis was performed using equipment purchased through NIH grant S10OD028632-01. We thank Dr. Zihua Wang for advice and a protocol for SMASH library preparation, Dr. Qing Gao for assistance with histological sample preparation and optimization, Jill Habel for assistance with mouse experiments. We are grateful for the support of Dr. Tawfiqui Bhuiya (Northwell Health) for tumor tissue sections and regret his untimely passing. We deeply appreciate the efforts of the Northwell Health Biobanking team for sample acquisition and the patients and their families who donated tissues for research. This research was supported by CSHL/Northwell Health support (D.L.S.), NCI 5P01CA013106-Project 3 (D.L.S.), Leidos Biomedical HHSN26100008 (D.A.T, D.L.S), and the Manhasset Women’s Coalition Against Breast Cancer (S.B.). This research was also supported by NCI Breast SPORE program P50-CA58223 (C.M.P), and U01CA238475-01 (C.M.P).

