## Supplementary files for "Patient-derived triple negative breast cancer organoids provide robust model systems that recapitulate tumor intrinsic characteristics"

**Supplementary Materials**

**Supplementary Table legends:**

**Table S1. Patient tissue information**: Clinical and culture properties of the 87 patient tumor tissues used in this study. Highlighted green and yellow samples are paired samples from two different patients.

**Table S2. CNA segments:** Final copy number segment file from SMASH data for all samples

**Table S3. RNA-seq sample information:** Sample details for samples used in transcriptomic profiling

**Table S4. RNA-seq signatures:** Gene expression signatures analysis for the various PDOs

**Table S5. Ki67 proliferation:** Percent Ki67 per organoid analysis for various normal and tumor PDOs

**Table S6. *In-vivo* transplant**: Overview of the PDO xenotransplant experiments N1 and N2, along with information on whether tumor formation and/or metastasis was observed.

**Table S7. Flow cytometry data and single cell experiment design:** Summary of EPCAM/CD49f flow cytometry data for the various lines. N= multiple passages for the tumor PDOs, and multiple different patient-derived normal organoid lines for the normal (NM0s) PDOs. Bottom panel: Experiment design for single cell experiment.

**Table S8. Differential gene expression of tumor vs normal LP cells**: Table of differential gene expression for comparing tumor LPs vs normal LPs

**Table S9. Cluster-specific gene markers:** Cluster specific markers for each of the clusters identified in TNBC PDOs

**Supplemental Figure Legends**

**Figure S1.** related to Figure 1,2 **A. & B.)** Variant allele frequencies comparisons on a patient-wise manner for the various SNVs identified in **A.)** tumor and/or PDOs **B.)** different passage PDOs. R represents Pearson correlation. **C.)** Oncoplot for pathogenic variants in tumor vs scraping derived organoids. **D.)** Oncoplot for tumor tissue (p0 organoids) for samples that did not result in successful organoid cultures. **E.**) Loss of copy number alterations in one PDO model overtime in culture.

**Figure S2.** related to Figure 3 **A.)** Heatmap of sample-by-sample distance matrix showing Euclidean distances between the different PDOs. Samples were clustered using hierarchical clustering with Ward linkage. type: Normal= reductive mammoplasty derived normal organoids, Paired Normal= Adjacent or Distal to the tumor paired normal, Normal outgrowth= no pathogenic mutations were found, Luminal= ER/PR+ organoids; driver mutation: Other= trace mutations (see Fig 1E), None= no pathogenic mutations were found, NA= not assessed **B.)** Copy number profiles of NH48N and NH48T **C.)** Gene expression signatures for selected modules. Pink box highlights the samples defined as LT-TNBCs **D.)** Gene expression signatures for luminal-progenitor gene sets and c-MYC amplification. Each dot represents a different PDO; Luminal N=12, Normal N= 7, TNBC N= 19, TNBC met= 4. **E.**) Copy number data for selected MYC genes

**Figure S3.** related to Figure 3 **A.)** Selected culture images for organoids that did not result in successful cultures. **B.)** Ki67 IHC and H&E images for PDOs derived from normal and tumor pairs of the same patient, scale bars= 100µm. **C.)** Distribution of percent Ki67

positive cells per organoid for the various normal and tumor-derived organoids. The numbers on top of violin points indicate the number of organoids counted. The black line in the violin represents median. Data is plotted using GraphPad Prism. **D.)** Growth rate of organoid formation as measure by 3D Cell Titre Glo assay d6 post seeding, bars represent average with standard error and individual data points are measurements at different passages per PDO. Data is plotted using GraphPad Prism. **E.)** Dose response curves for three different drugs on 5 different PDO lines, data-points are presented as mean$\pm$SEM and curve is the non-linear fit.

**Figure S4.** related to Figure 4 **A.)** H&E staining of PDO-X tumors from NH84T and NH93T that were not palpable and were only identified upon histological analysis. **B.)** Visualization of micro-metastasis and single cell metastasis using IHC for human mitochondria labeling in lymph nodes (upper panels) of the NOD/SCID mice transplanted with various PDOs. Bottom panels show human mitochondria labeling in lungs, liver and lymph-node from HCM-CSHL-0655-C50. Black arrows point to human mitochondria labeled micro- or single cell-metastasis. **C.)** CD117 and CK5/6 labeling of sections from an AdCC-like patient tumor (upper panels) and the PDO-X derived tumor (lower panels) from the same patient, scale bar on bottom panel=100$\mu$M.

**Figure S5.** related to Figure 5 **A.)** Flow-cytometry contour plots for EPCAM and CD49f for selected tumor PDOs (in red contours) overlayed onto normal-derived organoids (in blue contours). **B.**) Expression of ITGA6 and EPCAM from RNA-seq data for the different PDOs. Dark pink dots: true normal PDOs, light pink dots: paired normal PDOs and black dots: tumors. Circled region represents long term propagating organoids. **C.)** Different passage flow-cytometry plots for NH95T, NH85TSc and HCM-CSHL-0366-C50 showing stability of the cell populations over time in culture. **D.**) Flow-cytometry plots for some more TNBC and lymph-met derived organoids.

**Figure S6.** related to Figure 6 **A. & B.)** Cell cycle related **A.)** UMAP-plot and **B.)** gene expression in the cycling clusters **C.)** Cluster specific markers associated with each G1 cluster identified **D.)** Expression patterns of the various luminal/basal markers in normal and TNBC PDOs. HC-0366 is PDO HCM-CSHL-0366-C50 and HC-0655 is PDO HCM-CSHL-0655-C50 **E.)** Mammary lineage scores for each of the PDOs in specific clusters **F.)** Gene-set scores for each of the specified gene-set between the tumor and normal samples. **G.)** Motif enrichment for differentially expressed genes that were significantly upregulated in tumor LPs compared to normal LPs (clusters 2, 7 and 11). Score= -10*NES*padj.value **H.)** Representative culture images for three different PDOs treated with DAPT and MYCi **I.)** Normal organoid formation from sorted EPCAM+/CD49f+ and CD49f+ve populations from three different normal PDO lines in DAPT or MYCi containing medium. Significance was assessed by multiple-comparisons by two-way ANOVA ns= not significant, *** pvalue<0.005.

**Figure S7.** related to Figure 7 **A.)** Cluster distribution for each of the TNBC PDOs tested. **B.)** EPCAM and ITGA6 expression for all the TNBC samples **C.)** Feature plots showing mammary lineage scores for each of the PDOs in specific clusters **D.**) Boxplots showing scores for the different gene-sets tested **E.**) Feature plots showing the gene-set score distribution among the different TNBC PDOs. The first panel is the combined score distribution for all samples and provides the legend for the whole row. **F.**) Marker expression for one gene per cluster for the combined tumor organoids (first panel) and for individual TNBC organoids.

Fig S1

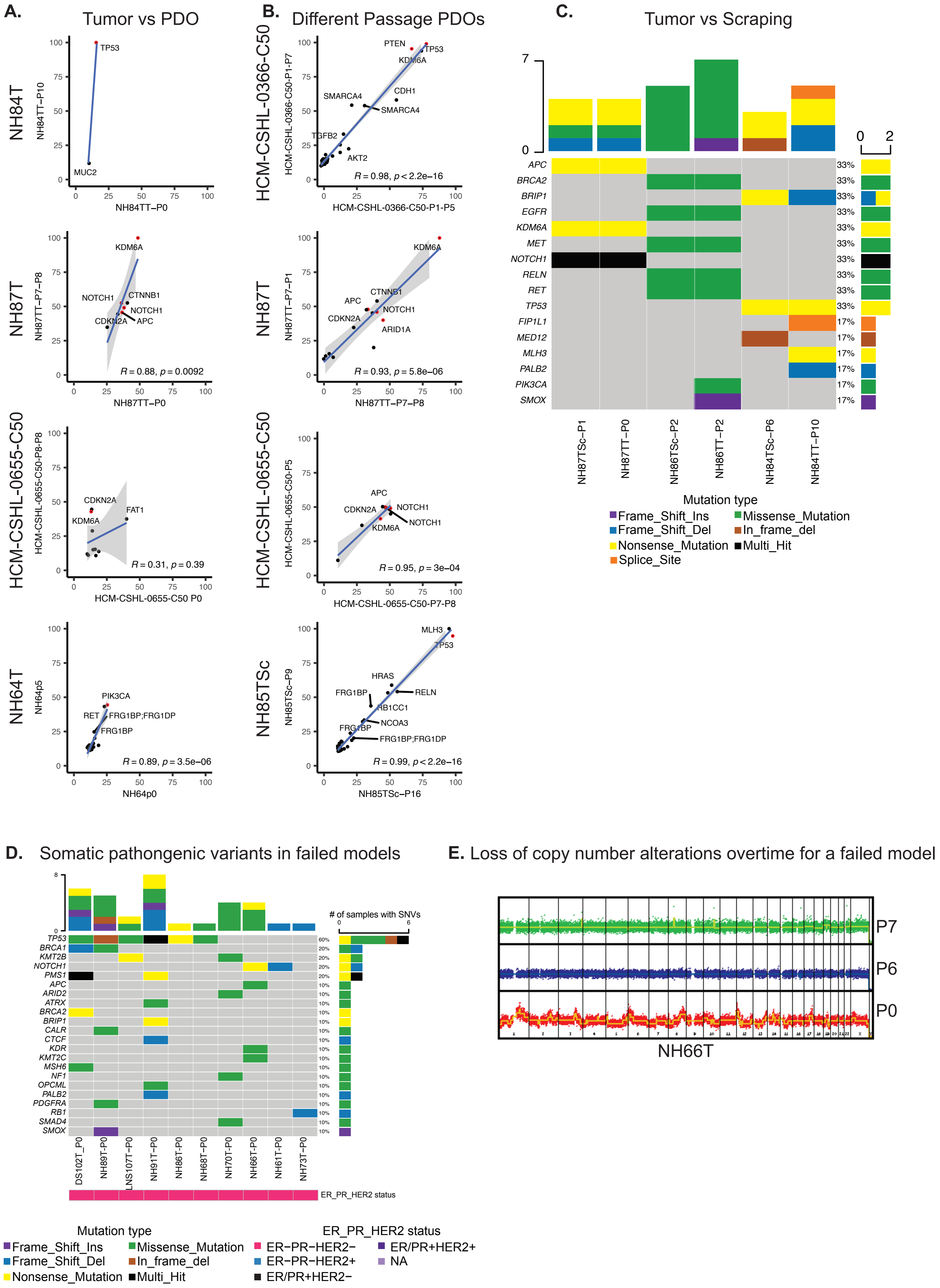

A.
Tumor vs PDO
NH84T
NH84TT–P10
NH84TT–P0
TP53
MUC2
NH87T
NH87TT–P7–P8
NH87TT–P0
KDM6A
NOTCH1
CTNNB1
NOTCH1
CDKN2A
APC
R = 0.88, p = 0.0092
HCM-CSHL-0655-C50
HCM-CSHL-0655-C50-P8-P8
HCM-CSHL-0655-C50 P0
CDKN2A
FAT1
KDM6A
R = 0.31, p = 0.39
NH64T
NH64p5
NH64p0
PIK3CA
RET
FRG1BP;FRG1DP
FRG1BP
R = 0.89, p = 3.5e–06
B.
Different Passage PDOs
HCM-CSHL-0366-C50
HCM-CSHL-0366-C50-P1-P7
HCM-CSHL-0366-C50-P1-P5
PTEN
TP53
KDM6A
CDH1
SMARCA4
SMARCA4
TGFB2
AKT2
R = 0.98, p < 2.2e–16
NH87T
NH87TT–P1
NH87TT–P7–P8
KDM6A
CTNNB1
APC
NOTCH1
CDKN2A
ARID1A
R = 0.93, p = 5.8e–06
HCM-CSHL-0655-C50
HCM-CSHL-0655-C50-P5
HCM-CSHL-0655-C50-P7-P8
APC
CDKN2A
NOTCH1
NOTCH1
KDM6A
R = 0.95, p = 3e–04
NH85TSc
NH85TSc–P9
NH85TSc–P16
MLH3
TP53
HRAS
FRG1BP
RELN
RB1CC1
FRG1BP
NCOA3
FRG1BP;FRG1DP
R = 0.99, p < 2.2e–16
C.
Tumor vs Scraping
APC
BRCA2
BRIP1
EGFR
KDM6A
MET
NOTCH1
RELN
RET
TP53
FIP1L1
MED12
MLH3
PALB2
PIK3CA
SMOX
NH87TSc–P1
NH87TT–P0
NH86TSc–P2
NH86TT–P2
NH84TSc–P6
NH84TT–P10
Mutation type
Frame_Shift_Ins
Missense_Mutation
Frame_Shift_Del
In_frame_del
Nonsense_Mutation
Multi_Hit
Splice_Site
D.
Somatic pathongenic variants in failed models
### of samples with SNVs
TP53
BRCA1
KMT2B
NOTCH1
PMS1
APC
ARID2
ATRX
BRCA2
BRIP1
CALR
CTCF
KDR
KMT2C
MSH6
NF1
OPCML
PALB2
PDGFRA
RB1
SMAD4
SMOX
DS102T_P0
NH89T–P0
LNS107T–P0
NH91T–P0
NH86T–P0
NH68T–P0
NH70T–P0
NH66T–P0
NH61T–P0
NH73T–P0
ER_PR_HER2 status
Mutation type
Frame_Shift_Ins
Missense_Mutation
Frame_Shift_Del
In_frame_del
Nonsense_Mutation
Multi_Hit
ER_PR_HER2 status
ER−PR−HER2−
ER/PR+HER2+
ER−PR−HER2+
NA
ER/PR+HER2−
E. Loss of copy number alterations overtime for a failed model
P7
P6
P0
NH66T

Fig S2

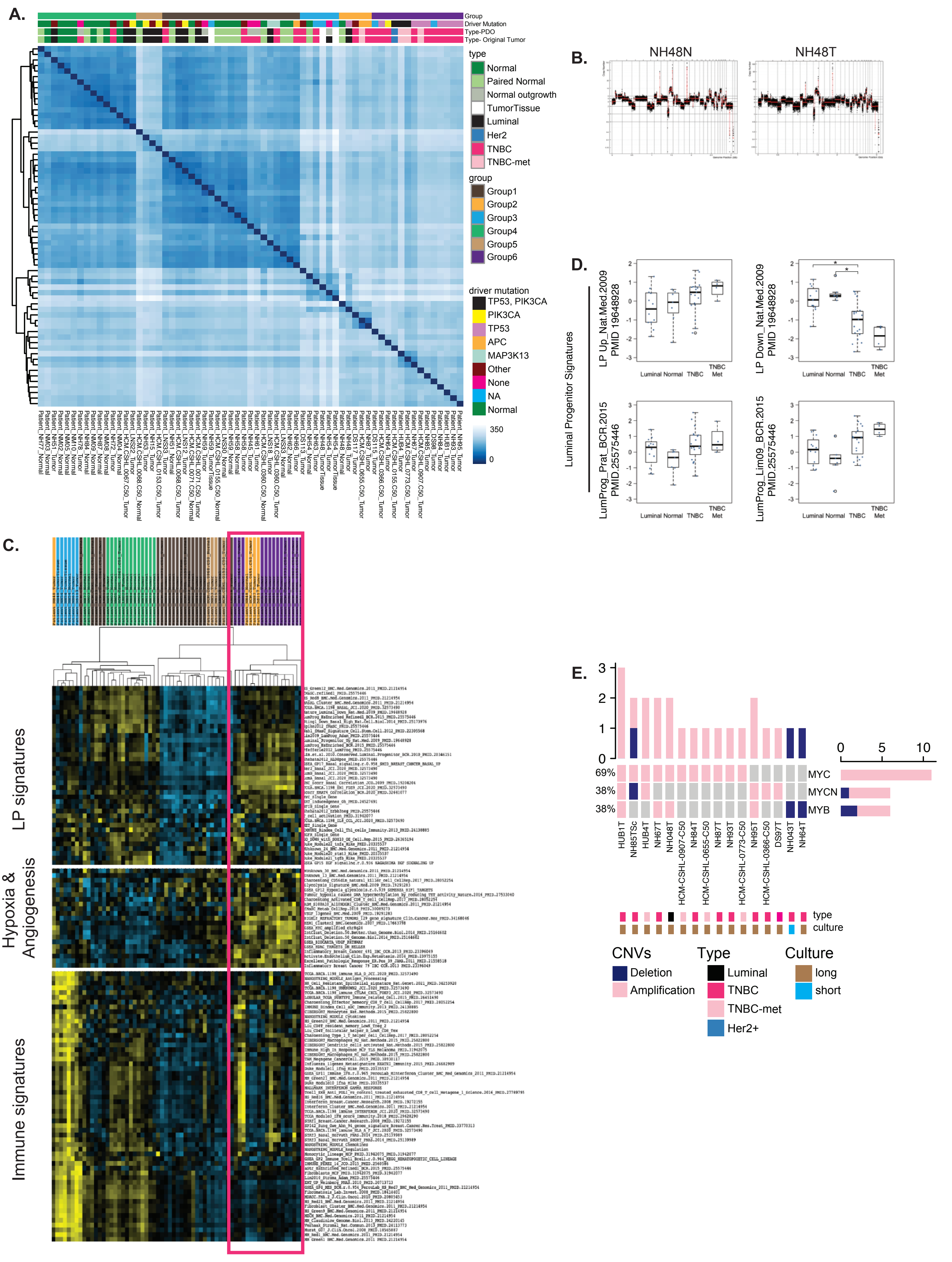
A.
Group
Driver Mutation
Type-PDO
Type- Original Tumor
type
Normal
Paired Normal
Normal outgrowth
TumorTissue
Luminal
Her2
TNBC
TNBC-met
group
Group1
Group2
Group3
Group4
Group5
Group6
driver mutation
TP53, PIK3CA
PIK3CA
TP53
APC
MAP3K13
Other
None
NA
Normal
350
0
B.
NH48N
NH48T
D.
Luminal Progenitor Signatures
LP Up_Nat.Med.2009
PMID 19648928
LP Down_Nat.Med.2009
PMID 19648928
LumProg_Prat_BCR.2015
PMID.25575446
LumProg_Lim09_BCR.2015
PMID.25575446
Luminal Normal TNBC TNBC Met
C.
LP signatures
Hypoxia & Angiogenesis
Immune signatures
E.
MYC
MYCN
MYB
69%
38%
38%
HUB1T
NH85TSc
HUB4T
NH67T
NH048T
HCM-CSHL-0907-C50
NH84T
HCM-CSHL-0655-C50
NH87T
NH93T
HCM-CSHL-0773-C50
NH95T
HCM-CSHL-0366-C50
DS97T
NH043T
NH64T
type
culture
CNVs
Deletion
Amplification
Type
Luminal
TNBC
TNBC-met
Her2+
Culture
long
short

Fig S3

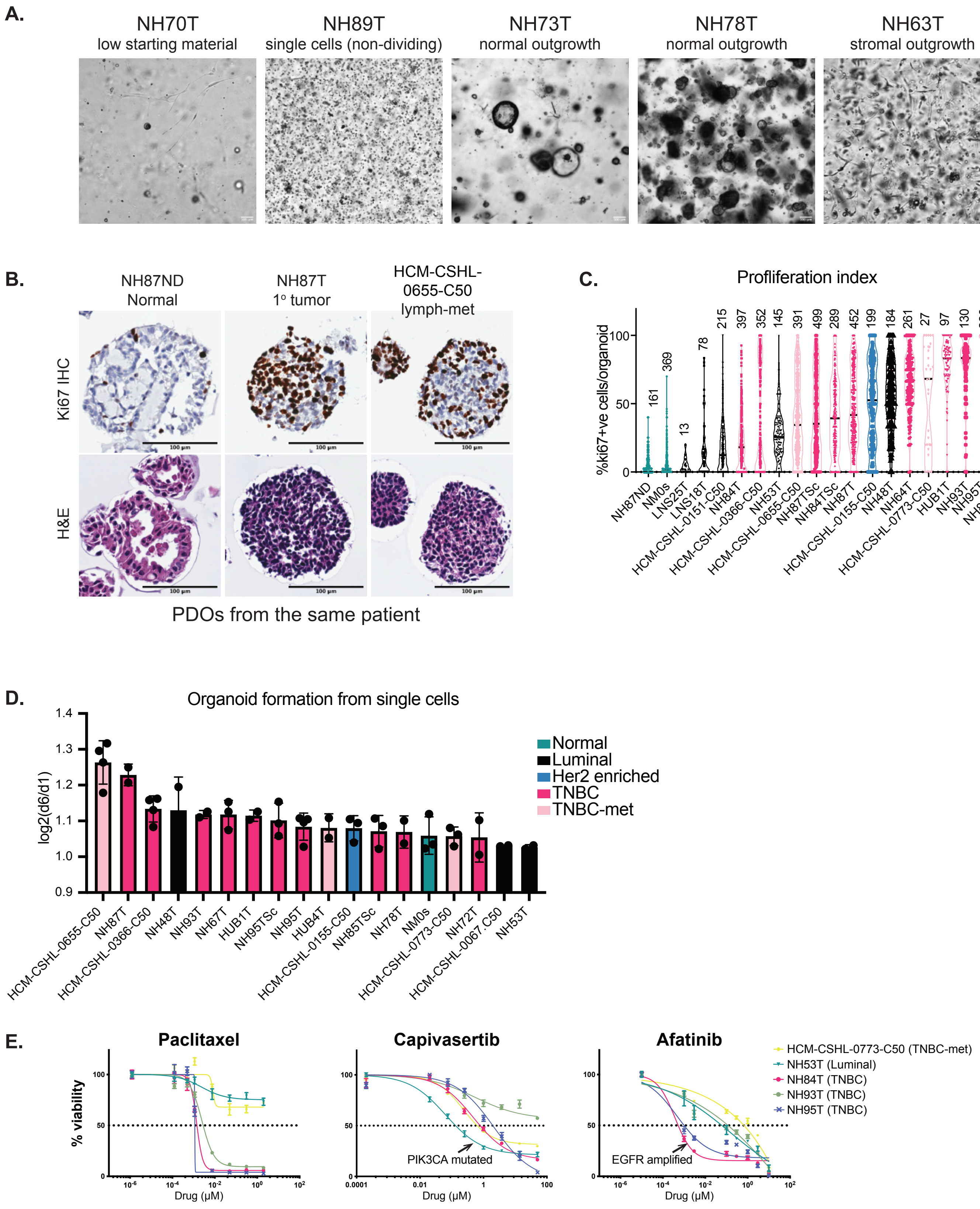
A.
NH70T
low starting material
NH89T
single cells (non-dividing)
NH73T
normal outgrowth
NH78T
normal outgrowth
NH63T
stromal outgrowth
B.
NH87ND
Normal
NH87T
1° tumor
HCM-CSHL-
0655-C50
lymph-met
Ki67 IHC
H&E
PDOs from the same patient
C.
Profliferation index
%ki67+ve cells/organoid
161
369
13
78
215
397
352
145
391
499
289
452
199
184
261
27
97
130
226
136
NH87ND
NM0s
LNS25T
LNS18T
HCM-CSHL-0151-C50
NH84T
HCM-CSHL-0366-C50
NH53T
HCM-CSHL-0655-C50
NH87TSc
NH84TSc
NH87T
HCM-CSHL-0155-C50
NH48T
NH64T
HCM-CSHL-0773-C50
HUB1T
NH93T
NH95T
NH85TSc
D.
Organoid formation from single cells
log2(d6/d1)
Normal
Luminal
Her2 enriched
TNBC
TNBC-met
HCM-CSHL-0655-C50
NH87T
HCM-CSHL-0366-C50
NH48T
NH93T
NH67T
HUB1T
NH95TSc
NH95T
HUB4T
HCM-CSHL-0155-C50
NH85TSc
NH78T
NM0s
HCM-CSHL-0773-C50
NH72T
HCM-CSHL-0067.C50
NH53T
E.
Paclitaxel
Capivasertib
Afatinib
% viability
Drug (µM)
PIK3CA mutated
EGFR amplified
HCM-CSHL-0773-C50 (TNBC-met)
NH53T (Luminal)
NH84T (TNBC)
NH93T (TNBC)
NH95T (TNBC)

Fig S4

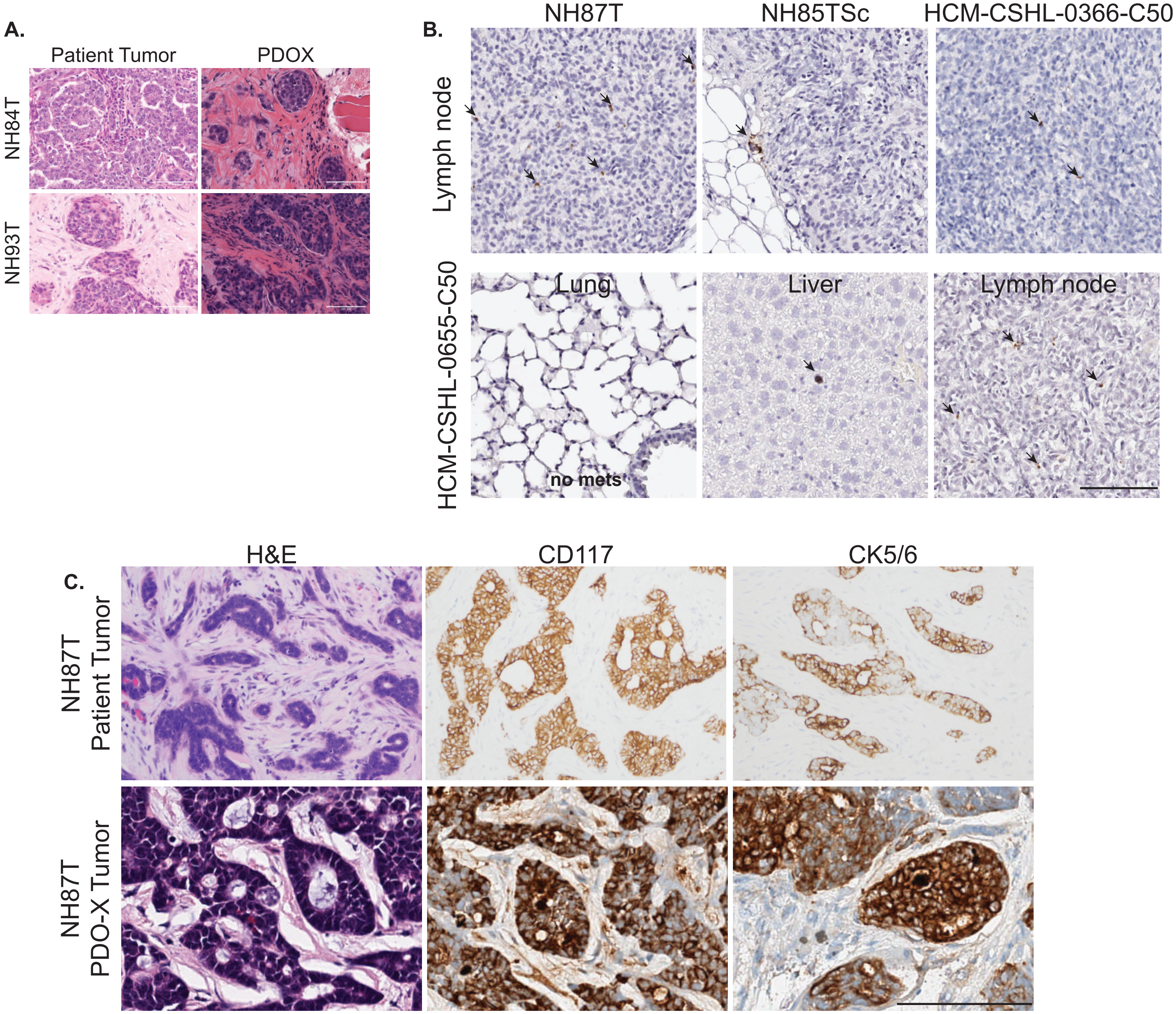

A.
Patient Tumor
PDOX
NH84T
NH93T
B.
NH87T
NH85TSc
HCM-CSHL-0366-C50
Lymph node
HCM-CSHL-0655-C50
Lung
Liver
Lymph node
no mets
C.
H&E
CD117
CK5/6
NH87T Patient Tumor
NH87T PDO-X Tumor

Fig S5

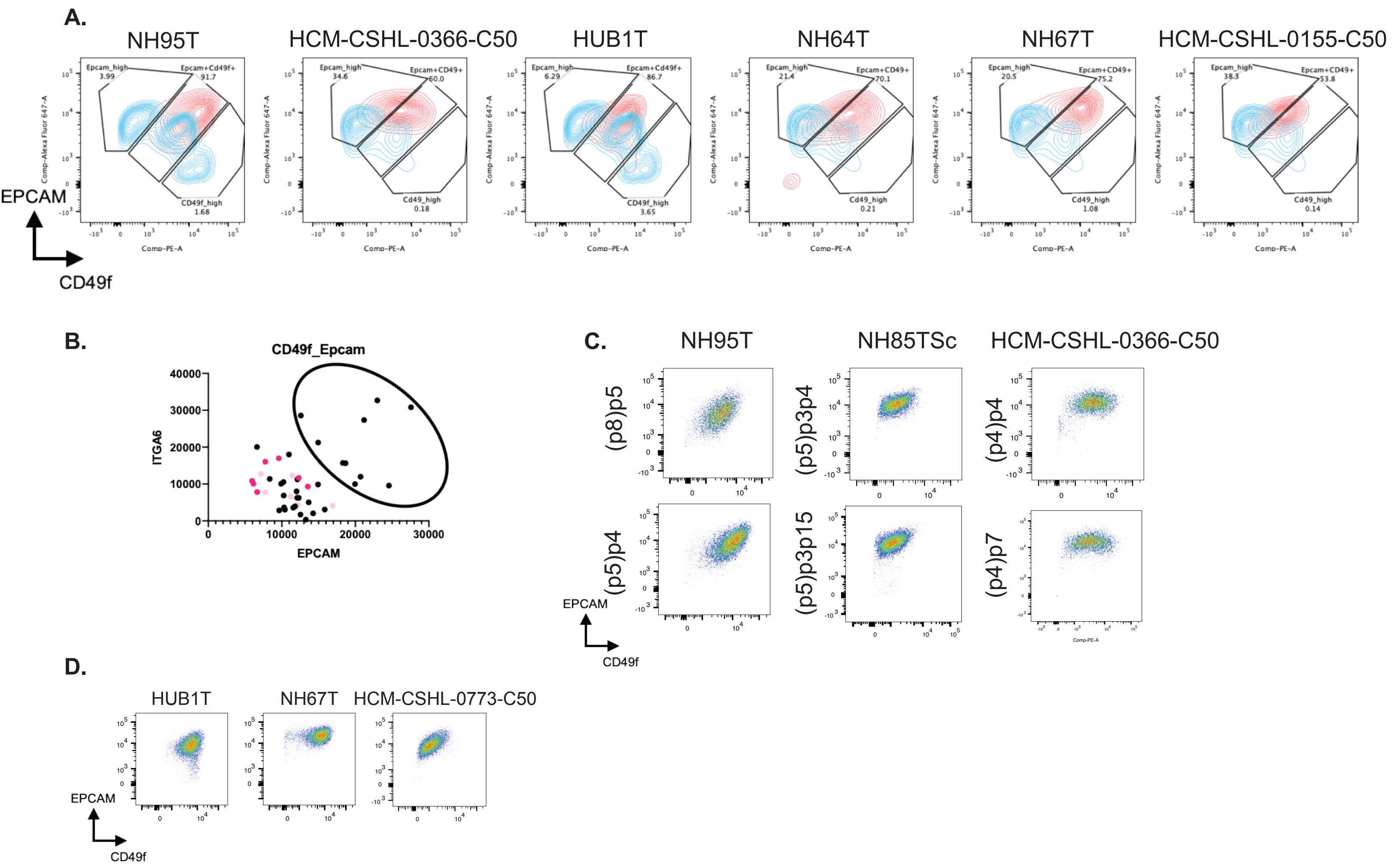
A.
NH95T
HCM-CSHL-0366-C50
HUB1T
NH64T
NH67T
HCM-CSHL-0155-C50
Epcam_high 3.99
Epcam+Cd49f+ 91.7
CD49f_high 1.68
Epcam_high 34.6
Epcam+CD49+ 60.0
Cd49_high 0.18
Epcam_high 6.29
Epcam+Cd49f+ 86.7
CD49f_high 3.65
Epcam_high 21.4
Epcam+CD49+ 70.1
Cd49_high 0.21
Epcam_high 20.5
Epcam+CD49+ 75.2
Cd49_high 1.08
Epcam_high 38.3
Epcam+CD49+ 53.8
Cd49_high 0.14
Comp-Alexa Fluor 647-A
Comp-PE-A
EPCAM
CD49f
B.
CD49f_Epcam
ITGA6
EPCAM
C.
NH95T
NH85TSc
HCM-CSHL-0366-C50
(p8)p5
(p5)p3p4
(p4)p4
(p5)p4
(p5)p3p15
(p4)p7
EPCAM
CD49f
D.
HUB1T
NH67T
HCM-CSHL-0773-C50
EPCAM
CD49f

Fig S6

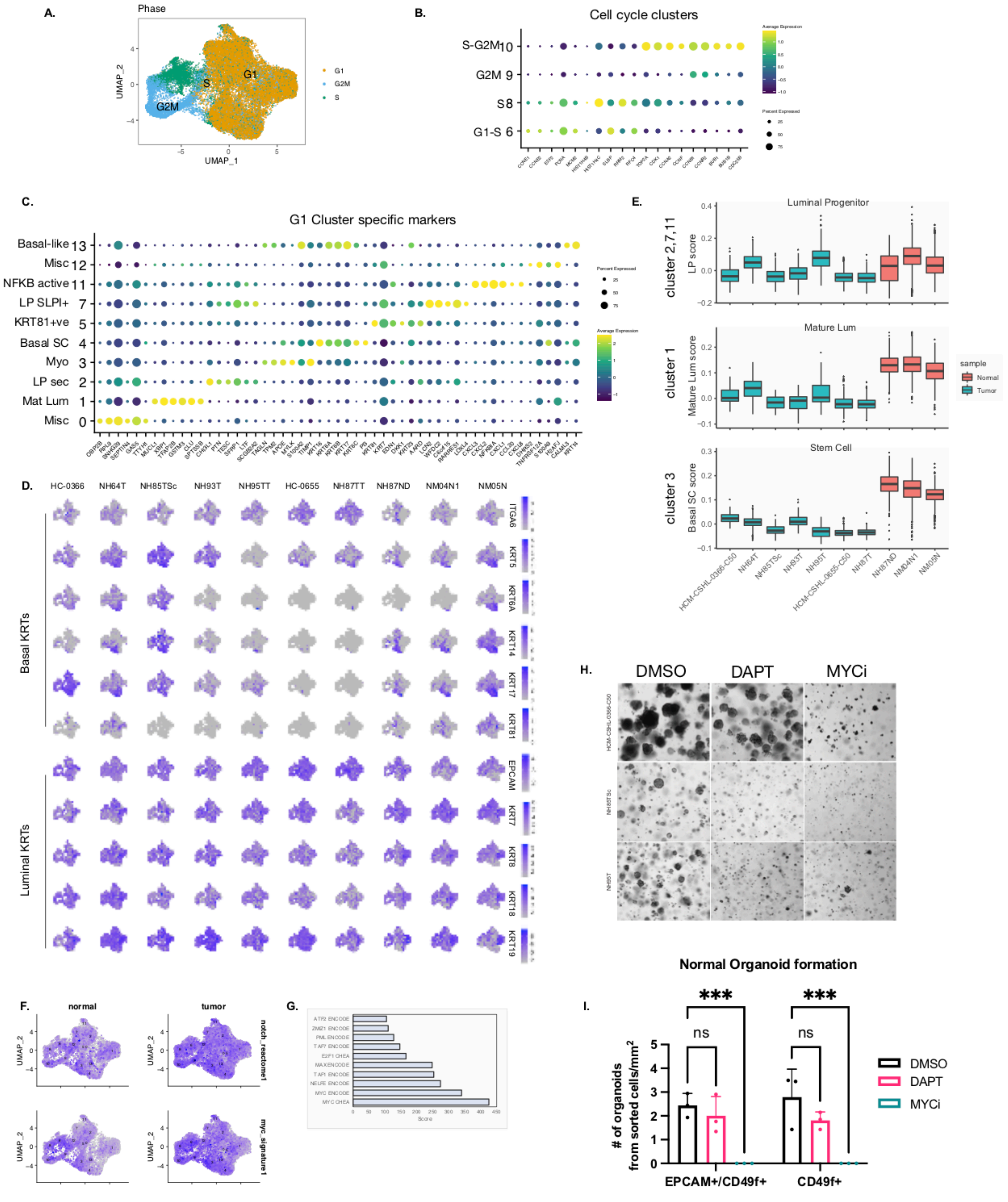
A.
Phase
G1
G2M
S
UMAP_1
UMAP_2
B.
Cell cycle clusters
S-G2M 10
G2M 9
S 8
G1-S 6
Average Expression
Percent Expressed
C.
G1 Cluster specific markers
Basal-like 13
Misc 12
NFKB active 11
LP SLPI+ 7
KRT81+ve 5
Basal SC 4
Myo 3
LP sec 2
Mat Lum 1
Misc 0
Percent Expressed
Average Expression
D.
HC-0366
NH64T
NH85TSc
NH93T
NH95TT
HC-0655
NH87TT
NH87ND
NM04N1
NM05N
ITGA6
KRT5
KRT6A
KRT14
KRT17
KRT81
EPCAM
KRT7
KRT8
KRT18
KRT19
Basal KRTs
Luminal KRTs
E.
Luminal Progenitor
cluster 2,7,11
LP score
Mature Lum
cluster 1
Mature Lum score
Stem Cell
cluster 3
Basal SC score
sample
Normal
Tumor
HCM-CSHL-0366-C50
NH64T
NH85TSc
NH93T
NH95T
HCM-CSHL-0655-C50
NH87T
NH87ND
NM04N1
NM05N
F.
normal
tumor
notch_reactome1
myc_signature1
G.
ATF2 ENCODE
ZMIZ1 ENCODE
PML ENCODE
TAF7 ENCODE
E2F1 CHEA
MAX ENCODE
TAF1 ENCODE
NELFE ENCODE
MYC ENCODE
MYC CHEA
Score
H.
DMSO
DAPT
MYCi
HCM-CSHL-0366-C50
NH85TSc
NH95T
I.
Normal Organoid formation
### of organoids from sorted cells/mm2
ns
***
EPCAM+/CD49f+
CD49f+
DMSO
DAPT
MYCi

Fig S7

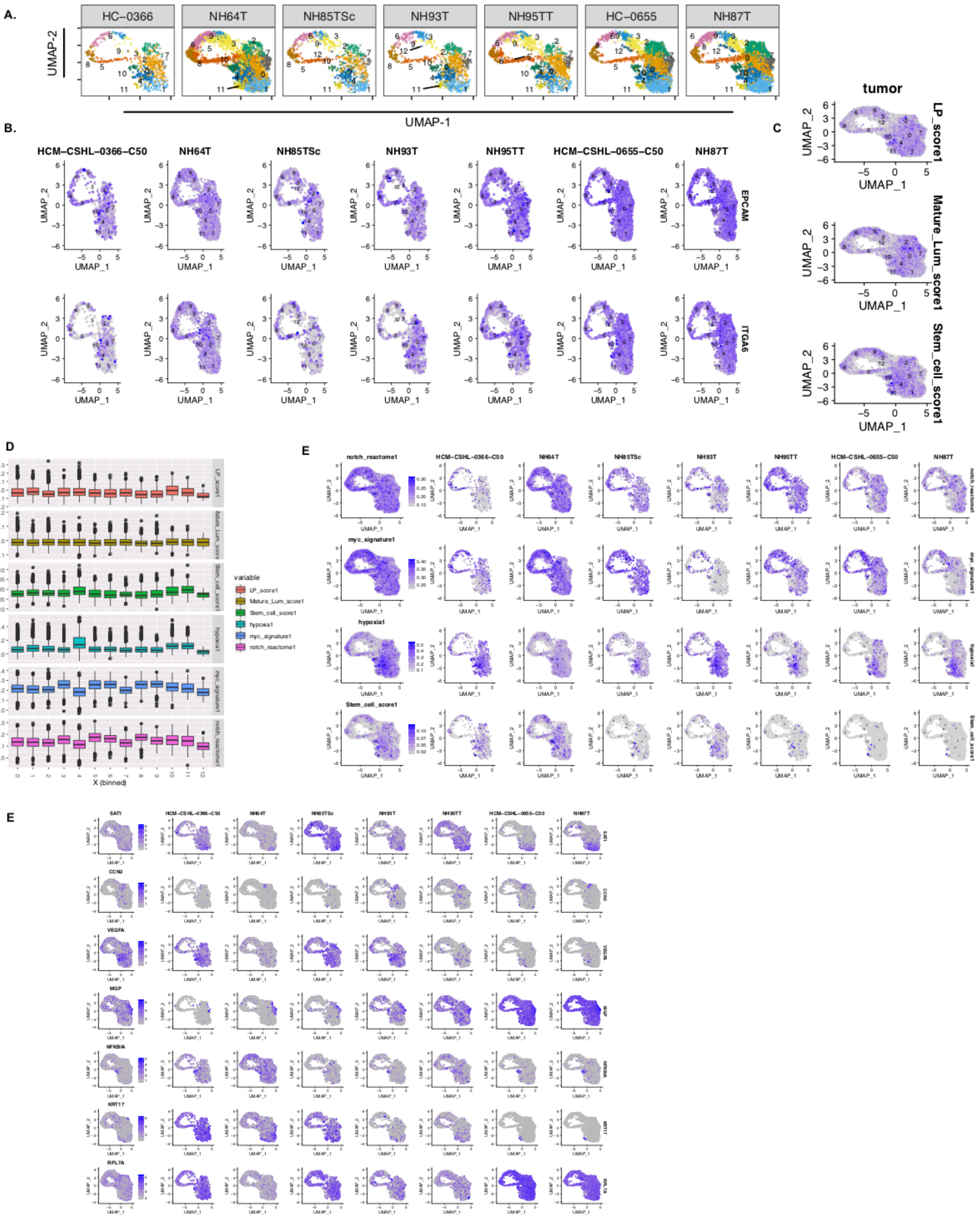
A.
HC−0366
NH64T
NH85TSc
NH93T
NH95TT
HC−0655
NH87T
UMAP-2
UMAP-1
B.
HCM−CSHL−0366−C50
NH64T
NH85TSc
NH93T
NH95TT
HCM−CSHL−0655−C50
NH87T
EPCAM
ITGA6
UMAP_1
UMAP_2
C
tumor
LP_score1
Mature_Lum_score1
Stem_cell_score1
D
variable
LP_score1
Mature_Lum_score1
Stem_cell_score1
hypoxia1
myc_signature1
notch_reactome1
value
X (binned)
E
notch_reactome1
myc_signature1
hypoxia1
Stem_cell_score1
E
SAT1
CCN2
VEGFA
MGP
NFKBIA
KRT17
RPL7A
